# Conserved Obg-type GTPases Sustain Protein Synthesis across Domains of Life

**DOI:** 10.64898/2026.05.13.724932

**Authors:** Sounak Saha, Christopher W. Hawk, Xin Chen, Aryaman Bhattacharya, Sarea Nizami, Melissa Pires-Alves, Gary Olsen, Hong Jin

**Affiliations:** Department of Biochemistry, University of Illinois at Urbana-Champaign, 600 S. Mathews Avenue, Urbana, IL 61801; Center for Biophysics and Quantitative Biology, University of Illinois at Urbana-Champaign, 600 S. Mathews Avenue, Urbana, IL 61801; Department of Microbiology, University of Illinois at Urbana-Champaign, 600 S. Mathews Avenue, Urbana, IL 61801; Carl R. Woese Institute for Genomic Biology, 1206 West Gregory Drive, University of Illinois at Urbana-Champaign, Urbana, IL 61801

**Keywords:** Protein synthesis, Ribosome, Ribosome Pause, GTPases, Ribosome-binding GTPase, Rbg, Developmentally-regulated GTP-binding protein, Obg

## Abstract

Translation is dynamically regulated to ensure proteins are synthesized at the correct time and location. Transient slowing or pausing of ribosomes enables coordination with cellular and environmental cues, yet how cells protect beneficial ribosomal pauses from premature activation of ribosome-associated quality control; and distinguish beneficial pauses from deleterious stalls, are not understood. Furthermore, sustaining productive protein synthesis under changing environments is fundamental to all cells. Here, we identify a conserved role for Obg-type ribosome-binding GTPases—Rbg protein in yeast and Drg proteins in humans—in promoting protein synthesis on halted ribosomes. We show that the essential bacterial Obg GTPase functionally rescues cell growth and translation in *Saccharomyces cerevisiae* and human cells lacking endogenous Rbg/Drg proteins, in a manner dependent on GTP hydrolysis. Consistently, Obg enhances puromycin incorporation into stalled ribosomes in a manner requiring GTP hydrolysis. Genome-wide mRNA-seq and proteomic analyses further reveal that Rbg1 associates with ribosomes translating proteins required for diverse cellular functions, including nuclear, mitochondrial, and secretory pathways. Together, these findings support a conserved mechanism in which Rbg/Drg GTPases protect physiological ribosomal pauses from premature engagement of stress and quality-control pathways, thereby sustaining productive translation across domains of life.

**Significance Statement:** How cells sustain protein synthesis in an ever-changing environment is fundamental to all cells. We identify a ribosome-binding GTPase—Rbg in yeast and Drg in humans—as a conserved regulator that protects physiological pauses and sustains protein synthesis. Remarkably, bacterial Obg functionally substitutes for eukaryotic Rbg/Drg in both yeast and human cells, revealing deep evolutionary conservation of this mechanism. Our data show that Obg-type GTPase promotes continued peptide-bond formation in a GTPase-dependent manner. Positioned near the ribosomal GTPase-associated center, these GTPases are also poised to compete with stringent-response and stress-activation factors that trigger translation shutdown. Together, the findings suggest that translation is controlled not only by canonical GTPases (such as EF-Tu and EF-G) that drive protein synthesis forward but also by Rbg/Drg GTPases that protect the catalytic cycle against premature arrest.

## Introduction

Proteins play critical functions in all biological processes, and all cellular proteins are synthesized in the ribosome using the nucleotide sequence of messenger RNAs (mRNAs) as a template. Protein synthesis is appreciated to be tightly regulated to meet the intrinsic and ever-changing demands of cell physiology and function. Decades of research have uncovered many molecular details of the translation process, including the intrinsic requirement that ribosomes briefly pause at each codon on the mRNA to carry out template-dependent translation. Beyond these natural periodic pauses, ribosomes do not translate the mRNA at a uniform speed (1, 2). They decelerate, temporarily halt, and may permanently stall over the course of translation. Translation of certain amino acids (e.g. proline, arginine, lysine) (3–5), limited availability of amino acids and charged tRNAs (6, 7), stalling sequences of the nascent peptides in the exit tunnel of the ribosome (8, 9), secondary structures in the mRNA (10), and interactions with RNA-binding proteins (11) are all demonstrated to modulate the speed of translation.

Despite the potential for attenuated protein output, a subset of translation pauses is programmed and functionally important. Rare codons are encoded in mRNA sequences to facilitate co-translational protein folding and modification (12–14), and sub-cellular localization (15). Translation pauses even allow cells to integrate multiple signals and coordinate different pathways in response to internal cues or stress (16). On the other end of the spectrum, a prolonged stall of the ribosome on an mRNA results in premature arrest of protein synthesis in the open reading frame, and elicits quality control mechanisms that recycle the stalled ribosomes and degrade the incomplete nascent polypeptide chain and troubled mRNAs (10, 17–19).

Since ribosomes move unidirectionally from the 5’ to 3’-end of a given mRNA, and mRNAs are frequently translated by multiple ribosomes simultaneously, extended ribosome pauses often lead to a collision of two or more translating ribosomes. Notably, cells undergoing fast proliferation and growth often produce an increased number of mature ribosomes. Even though translation initiation is rate-limiting, the elevated cellular ribosome concentration leads to increased ribosome flux on a given mRNA, thus increasing the likelihood of ribosome collisions. It was estimated that around 6-10% of translating ribosomes are in a collided state (20). Considering the sheer number of these events, it seems too costly for the cell to commit to quality control on every ribosome that is paused or collided with during translation.

A stochastic ribosome pause or collision, as an inherent feature of the translational process, is likely to be resolved naturally due to its transient nature. However, how does a cell distinguish between programmed, functionally beneficial pauses from problematic ribosome pauses or collisions? A programmed functionally beneficial ribosome pause or temporary collision, such as pauses that facilitate protein folding and localization, requires the translation machinery to recover and return to the translation pool. In contrast, a problematic ribosome pause and collision caused by faulty translational components requires activation of quality control mechanisms, such as ribosome-associated quality control (RQC), to disassemble the troubled translation machinery and to degrade the truncated protein (17, 18, 21). Furthermore, as ribosome pauses and collisions are nearly inevitable, how does the cell prevent premature or unnecessary activation of quality control pathways on a programmed ribosome pause? These are important questions that we set out to answer.

In this study, we report the evolutionarily conserved function of a ribosome-binding GTPase (Rbg in yeast, and so-called <u>d</u>evelopmentally <u>r</u>egulated <u>G</u>TP-binding (Drg) in human) in promoting protein synthesis in paused ribosomes. Rbg/Drg proteins are an ancient family of important but little understood GTPases that are ubiquitously expressed in actively growing and developing cells of plants, animals, and humans (3, 22). This family of proteins was named as such for a historical reason: Drg mRNA was highly expressed in neural precursor cells in developing mouse brains, but its expression decreased as their development progressed (23). Although the cellular function of Drg was unknown at the time of its discovery, it was presumed to be “developmentally regulated”, as its expression pattern followed those of many critical developmental proteins. After its discovery, Rbg/Drg mRNAs and proteins were found to be widely expressed at variable levels in cultured cell lines, as well as embryonic, postnatal, and adult tissues (24). An earlier phylogenetic study revealed that eukaryotes typically contain two Rbg/Drg genes, Drg1 and Drg2, whereas archaea only contain one (25), presumably an ancestral homolog of their eukaryotic counterparts. The amino acid sequences of eukaryotic Drg1 and Drg2 are highly homologous, containing a canonical G domain common to the GTPase Obg superfamily (26) (**Supplemental Figure 1**).

Eukaryotic Drg proteins are reported to heterodimerize with Dfrp proteins (<u>D</u>rg <u>f</u>amily regulatory <u>p</u>rotein) (27, 28), conferring stability to the Drg protein *in vivo* (27) and enhancing the GTPase activity of the Drg *in vitro* (29, 30). We discovered that yeast Rbg complexes suppress ribosome pauses and promote protein synthesis in paused ribosomes (3), demonstrating that this family of proteins functions at the crossroads of protein synthesis and quality control in eukaryotic cells. Here, we extend the functional conservation of Obg-type GTPase from the eukaryotic to the bacterial domain. We report that the bacterial ObgE GTPase, an essential protein in bacteria (26, 31), rescues the growth and translation defects in yeast and human cells caused by the deletion of their endogenous Rbg/Drg proteins. Furthermore, using an *in vitro* system devoid of ribosome biogenesis and RQC factors, we show that the Obg GTPase facilitates puromycin incorporation in stalled ribosomes in a GTPase-dependent manner (***when equimolar or 2:1 ObgE and stalled ribosomes are combined***). Results from combined selective mRNA sequencing and proteomic analysis (Rbg1-IP-mRNA seq and IP-LC-MS/MS experiments) show that Rbg1 targets ribosomes translating a broad range of proteins, many of which are critical to organelle functions and key cellular processes, suggesting a compromised cellular proteo-homeostasis when this GTPase is absent. Taken together, these results show that Rbg/Drg GTPases are physiologically important to helping protein biosynthesis in paused ribosomes, a function likely to be conserved from bacteria to humans.

## Results

### 1. Phylogenetic analysis reveals ancient conservation and early diversification of the Obg/Rbg/Drg GTPase family

To investigate the evolutionary relationships among Obg/Rbg-family GTPases, we constructed a protein-sequence phylogeny containing representative bacterial, archaeal, and eukaryotic homologs of Obg and Rbg (Figure 1). The resulting topology resolved two broadly distinguishable evolutionary groups corresponding to the Obg and Rbg families, while also revealing relationships between prokaryotic proteins and their eukaryotic descendants.

**Figure 1.**
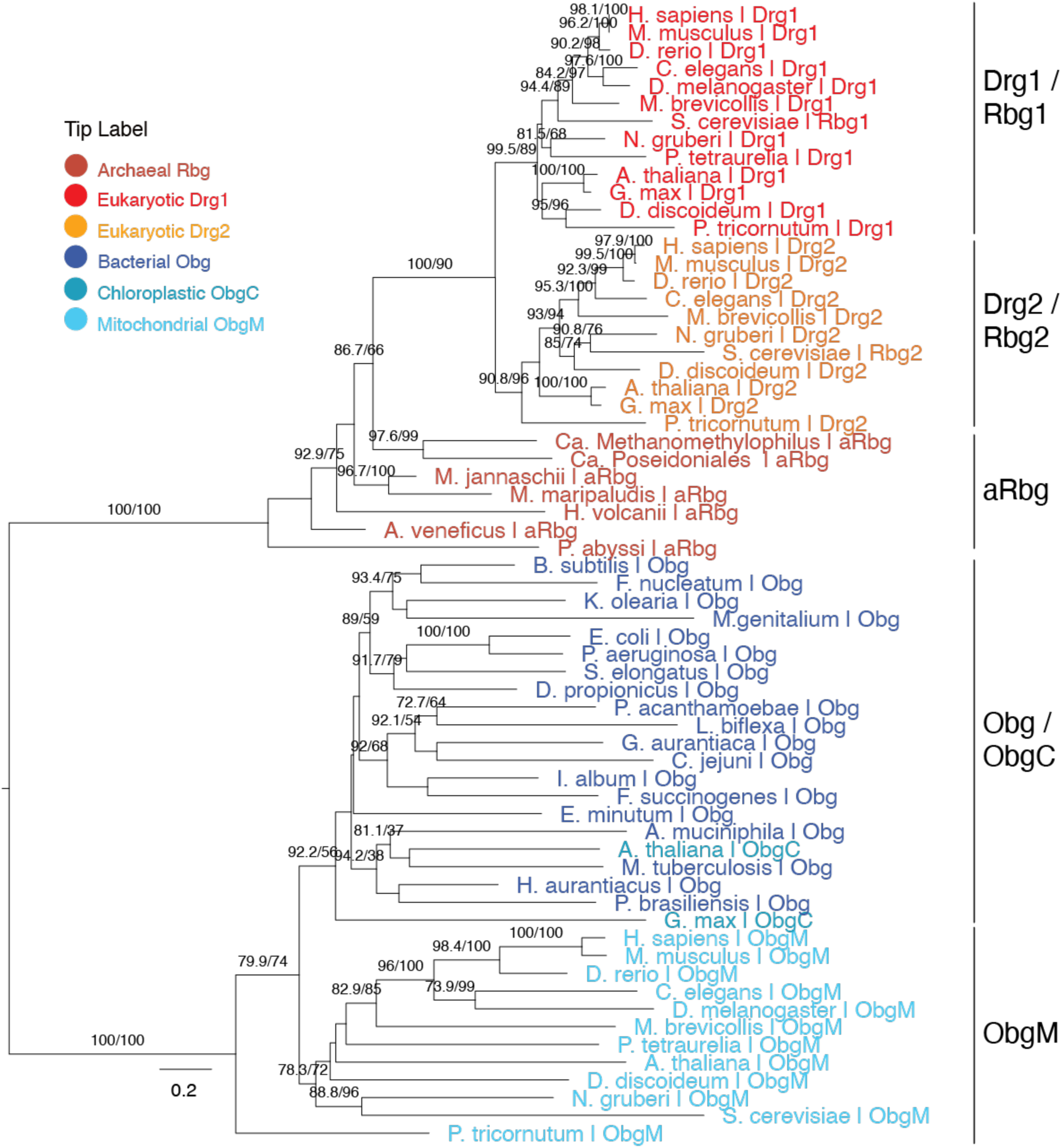
Phylogenetic relationships among Obg, and Rbg/Drg family GTPases across the three domains of life. The mid-point rooted phylogenetic tree was constructed from aligned protein sequences representing bacterial Obg proteins, archaeal and eukaryotic Rbg/Drg proteins, and eukaryotic organellar Obg homologs. Sequence labels specify the organism name and GTPase family. Rbg-family sequences are shown in red, orange and brown, while Obg-family sequences are shown in light blue, teal and dark blue. Numbers at internal nodes indicate branch-support values (SH-aLRT/UFBoot). Only SH-aLRT/UFBoot values greater than 70/70 are reflected. Branch lengths represent inferred evolutionary distances; the scale bar corresponds to 0.2 substitutions per site. Detailed domain or major taxonomic group, full organism name, and accession number are included in the Methods section.

Bacterial Obg proteins were distributed across a taxonomically diverse region of the tree. This group included homologs from Proteobacteria, Firmicutes, Actinobacteria, Cyanobacteria, Chloroflexota, Thermotogota, Spirochaetes, and several other bacterial phyla, consistent with the ancient and widespread conservation of Obg proteins in bacteria. Eukaryotic mitochondrial Obg proteins, designated ObgM, were positioned adjacent to bacterial Obg lineages rather than forming a completely separate lineage. Meanwhile, the plant chloroplast protein ObgC grouped among bacterial Obg homologs. These relationships are consistent with the bacterial ancestry of mitochondria and chloroplasts and suggest that organellar Obg proteins were inherited from the corresponding bacterial endosymbionts, followed by diversification within eukaryotic lineages.

Archaeal Rbg proteins (aRbg) and eukaryotic Rbg homologs formed the second major portion of the tree. Archaeal Rbg sequences were broadly distributed near eukaryotic Rbg branches, supporting an ancient relationship between the archaeal Rbg family and the eukaryotic Rbg proteins. Eukaryotic taxa generally encoded two related Rbg proteins, Rbg1 and Rbg2, and representatives of both lineages were detected across animals, fungi, plants, amoebozoans, alveolates, stramenopiles, and excavates. The presence of both paralogs in multiple diverged eukaryotic groups indicates that the Rbg1/Rbg2 duplication occurred early in eukaryotic evolution, before the diversification of the major extant eukaryotic supergroups.

Of note, we also carried out phylogenetic analysis of another distinct subfamily of Obg-type GTPase, Nog1, due to its well-known roles in ribosome biogenesis in eukaryotes. Obg/CgtA was known to perform a similar role in overseeing the maturation of pre-60S in the bacterial kingdom. Nog1 proteins formed a related but distinct evolutionary assemblage (**Supplemental Figure 2**). Eukaryotic Nog1 homologs from animals, fungi, plants, amoebozoans, stramenopiles, and excavates clustered together, with close relationships among orthologs from related species. Archaeal Nog1 homologs were positioned near the eukaryotic Nog1 lineage. This distribution indicates that Nog1 predates the radiation of modern eukaryotes and is consistent with its origin from an ancestral archaeal-type GTPase. The strong conservation of a single Nog1 lineage across diverse eukaryotic groups contrasts with the subsequent duplication and specialization observed for the Rbg proteins.

Several terminal relationships were strongly supported (SH-aLRT/UFBoot values both greater than 70), particularly those connecting closely related orthologs. By contrast, some internal branches separating ancient archaeal and eukaryotic Rbg sequences had relatively low support, preventing definitive reconstruction of their evolutionary relationship. The lack of complete monophyly for all sequences carrying the Rbg1 or Rbg2 annotation may reflect extensive divergence, differences in evolutionary rates, or uncertainty in assigning highly divergent homologs based on sequence alone. Nevertheless, the broad separation of Rbg proteins from Obg and Nog1, together with the occurrence of two Rbg paralogs across diverse eukaryotes, supports early functional diversification within this conserved GTPase superfamily.

Collectively, the phylogeny supports a model in which bacterial Obg, archaeal Rbg/Nog1, and eukaryotic Rbg-related proteins arose from an ancient family of P-loop GTPases. Eukaryotes retained specialized Nog1 and duplicated Rbg1/Rbg2 lineages, while bacterial-derived Obg homologs were preserved in mitochondria and chloroplasts. These findings suggest the deep evolutionary conservation of the Obg/Rbg pathway while revealing lineage-specific expansion and specialization during eukaryotic evolution.

### 2. Bacterial Obg rescues Δrbg1 yeast cell growth in the presence of anisomycin

The evolutionary persistence of the Obg-type GTPase family suggests that its core biochemical activity performs a fundamental cellular function. Moreover, the relationship between bacterial Obg proteins, archaeal Rbg proteins, and eukaryotic Rbg/Drg proteins provided the rationale to test whether their evolutionary conservation extends beyond sequence and structure to biological function. We therefore next examined whether Obg/Rbg/Drg proteins from evolutionarily distant organisms could functionally substitute for one another using cellular and biochemical assays.

To determine whether the cellular function of Drg/Rbg/Obg proteins is likewise conserved across the three domains of life, we studied cell growth behavior in the presence of the bacterially-derived antibiotic anisomycin, which stalls protein biosynthesis by inhibiting peptidyl transferase activity of the ribosome (32, 33). *S. cerevisiae* cells lacking their native Rbg1 are observably less viable in the presence of anisomycin when compared to wild-type (WT) cells (3). In the initial phase of the investigation, we expressed yeast (*S. cerevisiae*), bacterial (*E. coli*), and archaeal (*Methanosarcina acetivorans)*, namely Rbg1, ObgE, and aDrg, on a constitutively active PGK promoter via pJK372 plasmid in *Δrbg1* yeast cells lacking the endogenous rbg1 gene and assayed their growth using a spot assay. As expected, when compared to WT, *Δrbg1 S. cerevisiae* exhibited significantly reduced cellular viability when grown in the presence of 10 and 12.5 µg/mL anisomycin (**Figure 2A**, lines 1 and 2). However, supplementing *Δrbg1* yeast with Rbg1, archaeal aDrg, and ObgE proteins (**Figure 2A**, line 3, line 5 and line 6, respectively) visibly augments cellular growth compared to the *Δrbg1* yeast cell transformed with the same pJK372-PGK plasmid lacking a protein-coding sequence (empty vector, abbreviated as EV). Plasmid-expressed *E. coli* EF-Tu and a GTPase-deficient S79N mutant of Rbg1 show limited or no rescue of the slowed growth phenotype caused by Rbg1 deletion (**Figure 2A**, line 4 and line 7, respectively). Biological replicates of spot assays are reported in **Supplemental Figure 3A.**

**Figure 2.**
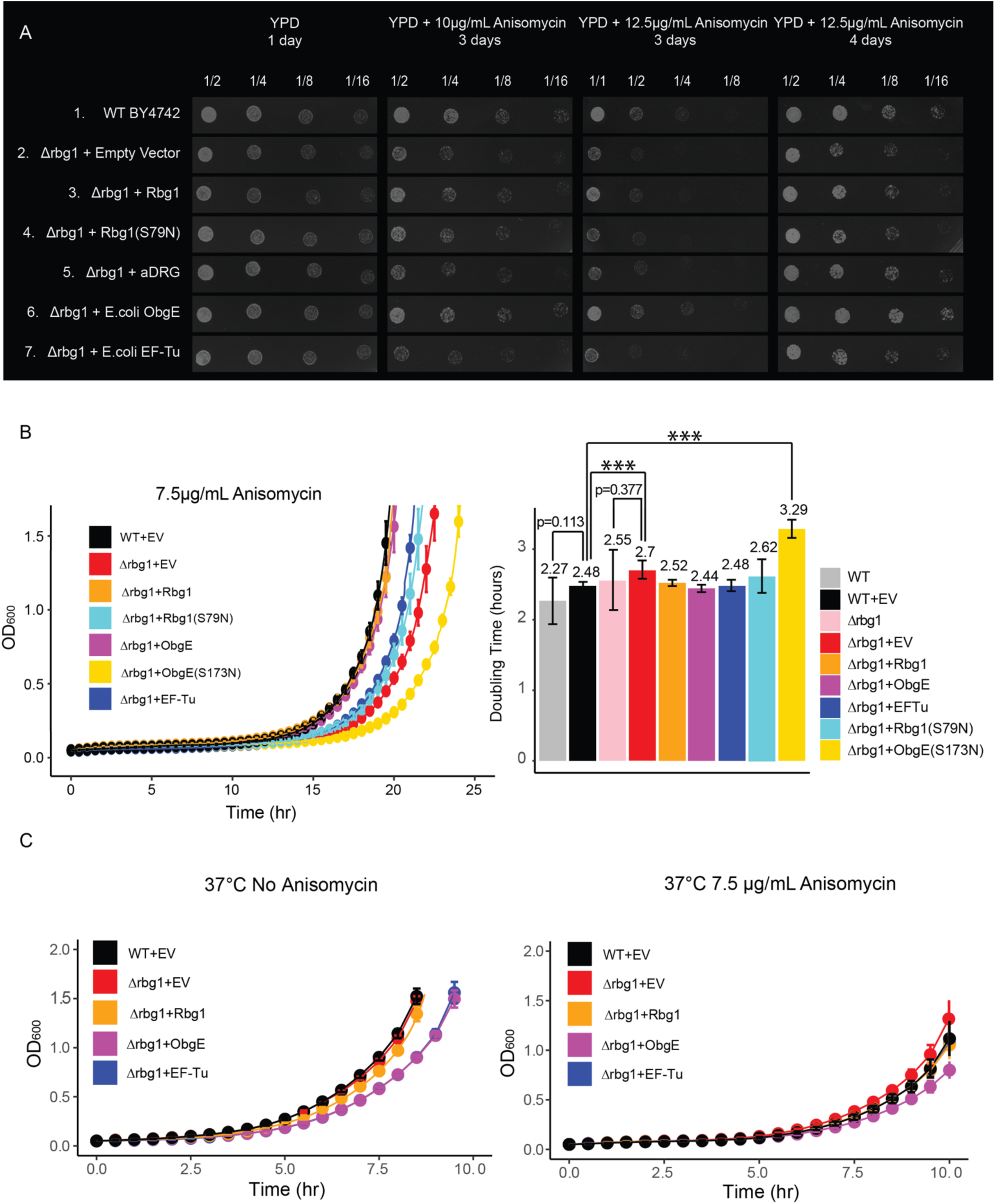
Bacterial Obg proteins compensate for lost GTPase-dependent Rbg function in yeast. **A. Bacterial Obg and archaeal Drg proteins compensate for the lost Rbg1 function in yeast cells.** Serial dilutions of wild type and *Δrbg1 S. cerevisiae* that were transformed with the indicated vectors were spotted on YPD plates with 0 µg/mL, 10 µg/mL, and 12.5 µg/mL anisomycin. Vector-derived yeast Rbg1 expression restores cell viability to near-WT levels and cells expressing bacterial ObgE and archaeal Drg (aDrg) exhibit higher cellular growth than cells supplemented with the native yeast Rbg1. **B. Plasmid-expressed Rbg1 and ObgE rescue *Δrbg1* yeast growth.** WT-like growth is observed in strains expressing Rbg1 or ObgE, while GTPase-deficient mutants Rbg1(S79N) and ObgE (S173N) fail to rescue growth in 7.5 µg/mL anisomycin. A bar plot shows that ObgE and EF-Tu yield the fastest log-phase doubling times among the expression constructs tested. Data points represent mean A_600_ values from nine replicates at the indicated time points. Error bars indicate the standard error of the mean. Doubling times are indicated as numbers above each bar. **C. Native Rbg1 is required for yeast cells under stress.** Growth curves of WT and *Δrbg1* strains with plasmid-based complementation, monitored by optical density at OD_600_ over 10 hours at 37 °C. Strains carried either an empty vector (EV) or a vector expressing the indicated protein (Rbg1, ObgE, or EF-Tu). Growth at 37 °C without anisomycin (Left panel), and growth at 37°C with 7.5 μg/mL anisomycin. The *Δrbg1* deletion strain exhibits a growth defect relative to WT under both conditions, which is rescued by re-expression of Rbg1 but not by the heterologous GTPases ObgE or EF-Tu. Note that in the absence of anisomycin, WT and Δrbg1 cells with only an empty vector, ie, WT+EV and Δrbg1+EV, grow similarly; these cells grow comparably, or only slightly better than the Δrbg1+Rbg1 cell at 37 °C. In the presence of anisomycin, WT cells with only an empty vector (WT+EV) display enhanced growth relative to all *Δrbg1*-derived strains, indicating that the presence of a suitable amount of Rbg1 contributes to translational robustness under stress conditions. Data points represent the mean ± SEM of 8 biological replicates.

To gain a more quantitative understanding of these cellular growth patterns, we monitored their growth over time in liquid medium. First, we assayed whether transformation with the pJK372-PGK plasmid caused any cell growth changes. Non-transformed cells entered log-phase growth earlier than those transformed with the empty vector, likely due to the selective media used for the initial selection of the cells harboring the transformed plasmids. However, all four cells, WT, WT+EV, *Δrbg1, and Δrbg1* + EV, have similar log-phase doubling times (**Supplemental Figure 3B**). Additionally, since an excess amount of Rbg1 in the cell is inhibitory based on the structural evidence (3), to see the relative protein abundance between native and pJK372-PGK plasmid-expressed Rbg1, we measured steady-state Rbg1 protein levels in WT, *Δrbg1+*Rbg1, and *Δrbg1*+Rbg1(S79N) via immunoblotting. As shown in **Supplemental Figure 3C**, Rbg1 expressed under the PGK promoter shows an approximately 2-fold increase in cellular concentration compared to chromosomally expressed Rbg1.

Since the presence of the empty vector affects lag-to-log phase transition in the growth behavior (**Figure 2A** and **Supplemental Figure 3B**), we proceeded with the WT and Δ*rbg1* yeast cells transformed with the empty vector (WT+EV and Δ*rbg1*+EV) as the controls for our liquid growth assays. As shown in **Supplemental Figure 3D**, ObgE– and Rbg1–supplemented *Δrbg1* cells grow similarly to the WT+EV and Δ*rbg1*+EV controls when grown in the absence of, or at low concentrations of anisomycin. However, differences between their growth rates became visible when the concentrations of anisomycin increased, and expression of plasmid-derived ObgE and Rbg1 rescued the growth defect resulting from yeast Rbg1 deletion. Of note, the key difference in cell growth between the Δ*rbg1* versus WT, as well as Obg– or Rbg1-supplemented Δrbg1 cells is that ***a longer lag phase*** is observed when Rbg1/Obg GTPase is absent. The physiological significance of the delayed lag-to-exponential transition in the cell growth phase in the absence of Obg/Rbg GTPase will be discussed in more detail in the Discussion section.

When assayed in YPD containing 7.5 µg/mL anisomycin, we observed similar rescue of the cell growth defect when *Δrbg1* cells were supplemented with WT Rbg1 and ObgE. As shown in **Figure 2B**, among the cells tested, *we observed earlier entry into exponential-phase growth in Δrbg1 cells supplemented with ObgE and Rbg1 proteins, a growth behavior similar to the WT+EV controls.* Both *Δrbg1* cells expressing ObgE and Rbg1 showed similar exponential phase doubling times as that of the WT+EV control, and all three were faster than the Δ*rbg1*+EV control, where none of the GTPases are supplemented.

Of note, under this experimental condition, *Δrbg1* cells expressing *E. coli* EF-Tu show relatively slower exponential phase entry, but similar doubling times to the WT+EV control. This result is likely due to a heterologous interaction between *E. coli* EF-Tu and the yeast 80S ribosome that decreases the GTP-hydrolysis rate (kcat) in EF-Tu as well as EF-Tu’s dissociation from the 80S ribosome after GTP hydrolysis. Additionally, Rbg1 and EF-Tu are structurally similar translational GTPases. Clearly, more investigations are required to uncover the underlying molecular mechanisms of the observed phenomenon on EF-Tu.

It’s important to note that faster growth under standard laboratory conditions does not necessarily imply an overall fitness of an organism. When the same experiments are carried out at an elevated growth temperature (37 °C) or in the presence of the same amount of anisomycin at 37 °C, supplementation of ObgE cannot rescue the growth as well as the native Rbg1 supplementation in the *Δrbg1* cell (**Figure 2C**). As expected, WT cells grow the best under those stress conditions (**Supplemental Figure 3E**). This phenomenon also indicates that the observed growth defect is specific to the presence of anisomycin which is known to bind to the PTC of the ribosome to stall translation.

Rbg/Drg homologs from all species share the conserved G domain comprised of well-known G1-G5 motifs. A conserved serine residue of the G1 motif (S79 in yeast Rbg1) was shown to be critical for protein’s cellular function, where the S79N mutation diminishes Rbg1 function in rescuing cell growth under the *Δrbg1Δrbg2Δslh1* (34). A related mutant Rbg1, containing alanine substitutions, including at the S79 residue, in the G1 motif (G77A, K78A and S79A) reduces the association of the Rbg1/Tma46 complex with translating ribosomes (34). It is important to note that S79 is located at the tip of the alpha helix in the G1 motif that is critical for nucleotide-binding. In our experiment, the S79N mutant of the native yeast Rbg1 cannot restore cell viability to the level of the WT protein in the spot assay (**Figure 2A, Line 4).** This mutant also grows more slowly than wild-type Rbg1-supplemented *Δrbg1* cells in liquid cultures (**Figure 2B**, light blue line). Furthermore, an analogous mutant in ObgE, S173N, abolishes its ability to rescue the lost endogenous Rbg1 function (**Figure 2B**, yellow line). These observations suggest that the nucleotide-binding and GTPase activity of the Rbg/Obg is critical to its cellular function. In this case, deficiencies in nucleotide-binding, thereby a lack of GTP hydrolysis, by Rbg/Obg GTPases may affect their ribosomal interaction and the protein’s function on the ribosome, as we will discuss in the Discussion section. Taken together, data from these straightforward experiments demonstrate a distinct and conserved GTPase-dependent cellular function shared among the Obg/Rbg proteins from bacteria to eukaryotes.

### 3. Bacterial ObgE GTPase helps restore translation through a polyarginine stall sequence in Rbg1-deficient yeast

Is the conserved cellular Rbg/Drg function that is evident from the anisomycin growth assay related to cytoplasmic protein biosynthesis? We noted from the onset of this investigation that bacterial ObgE directly binds to eukaryotic ribosomes (**Supplemental Figure 4**). To further investigate whether the observed rescue reflects a specific effect on translation rather than a nonspecific improvement in growth, we examined the phenotype using a translation reporter assay. Unlike growth-based measurements, this reporter directly monitors translation-dependent activity, and therefore provides a more specific readout of ribosome function.

We previously showed that yeast Rbg proteins promote efficient translation when ribosomes decelerate or pause on mRNAs. Using a standard R12 stalling reporter, the presence of yeast Rbg1 enhances translation and mRNA stability (3). To test whether the translation-promoting function of Obg-type GTPases is conserved from bacteria to single-cell eukaryotes, we used a dual-fluorescence reporter containing mCherry upstream and eGFP downstream of 12 consecutive arginine codons (R12; CGX), which create a strong translational impediment (**Figure 3A**). Because mCherry is synthesized before the R12 sequence whereas eGFP production requires ribosomes to translate through it, the eGFP/mCherry fluorescence ratio provides a measure of productive translation beyond the polyarginine sequence.

**Figure 3.**
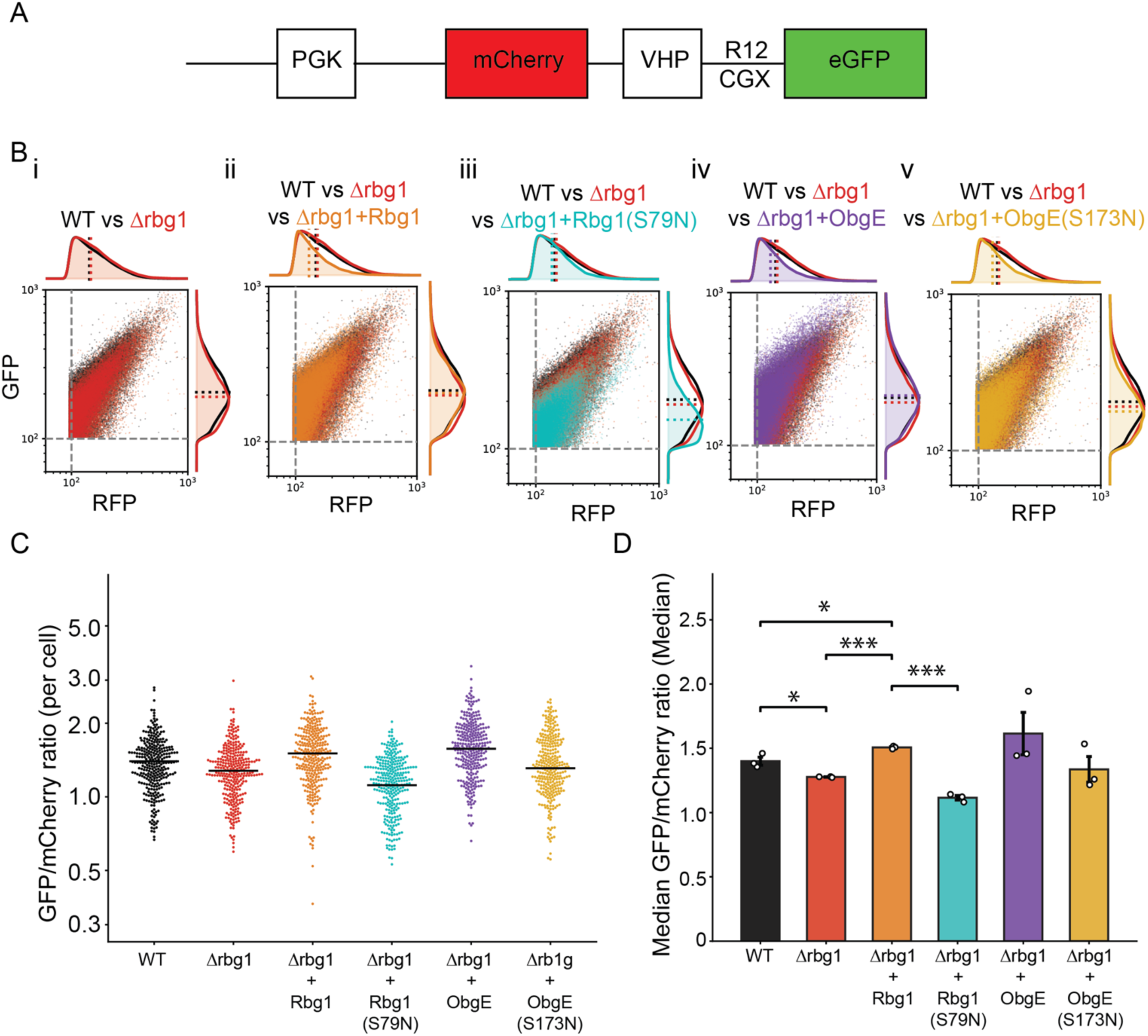
Bacterial ObgE promotes translation through a polyarginine sequence in Rbg1-deficient yeast. **A. Schematic of the dual-fluorescence reporter expressed from the PGK promoter.** The reporter encodes mCherry upstream and eGFP downstream of a VHP linker and 12 consecutive arginine codons (R12; CGX). Because eGFP synthesis requires translation through the R12 sequence, the eGFP/mCherry ratio reports productive downstream translation. **B. Representative flow-cytometry profiles comparing wild-type (WT) and Δrbg1 yeast with Δrbg1 cells expressing bacterial ObgE, GTPase-defective ObgE(S173N), GTPase-defective yeast Rbg1(S79N), or wild-type Rbg1.** Flow cytometry scatter plots show mCherry and eGFP fluorescence in individual cells, with corresponding marginal density distributions. Grey dashed lines indicate the 10² RFU gating threshold applied to both channels; marginal traces show kernel density estimates of RFP (top) and GFP (right), with dotted lines marking distribution medians. WT (black) and Δrbg1 (red) carry the *LEU2*-marked reporter plasmid alone, reference strains are overlaid in every panel. Each subsequent panel additionally shows one Δrbg1 strain carrying a second plasmid — identical in backbone to the reporter plasmid but marker-swapped to *URA3* — expressing the bacterial ortholog ObgE (purple) or its P-loop mutant ObgE(S173N) (gold), or the yeast Obg-family GTPase Rbg1 (orange) or its P-loop mutant Rbg1(S79N) (cyan). **C. Distributions of the eGFP/mCherry ratio among individual cells under the indicated conditions.** Single-cell distributions of the eGFP/mCherry ratio for each condition, shown as beeswarm plots on a log-scaled axis. For visual clarity, 300 cells per condition were randomly subsampled from the pooled events described in (B); horizontal black bars indicate the per-condition median of the full pooled population. **D. Quantification of the median eGFP/mCherry ratio from three independent biological replicates.** Open circles represent individual biological replicates; bars show the mean of the replicate medians, and error bars indicate SEM. Loss of Rbg1 modestly but significantly reduced downstream eGFP synthesis. Expression of either wild-type bacterial ObgE or yeast Rbg1 increased the reporter ratio, whereas the GTPase-defective Rbg1(S79N), ObgE(S173N) mutants failed to rescue the defect. Statistical significance was determined using an unpaired two-tailed t-test; *P* < 0.05 and \*\**P* < 0.001.

Flow-cytometric analysis revealed a modest but statistically significant reduction in the eGFP/mCherry ratio in *Δrbg1* cells compared with wild-type cells, indicating that loss of Rbg1 impairs translation through the R12 sequence (**Figure 3B, i, and Figure 3C and 3D**). Expression of wild-type yeast Rbg1 in *Δrbg1* cells increased the reporter ratio and restored translation beyond the polyarginine sequence (**Figure 3B, ii, and Figure 3C and 3D**). By contrast, the GTPase-defective Rbg1(S79N) mutant ailed to rescue the defect and produced a significantly lower reporter ratio than wild-type Rbg1 (**Figure 3B, iii, and Figure 3C and 3D**). These results indicate that Rbg1 promotes productive translation through the R12 sequence and that this activity depends on its functional GTPase domain.

Remarkably, expression of bacterial ObgE in Δrbg1 yeast also increased the eGFP/mCherry ratio, producing a reporter signal comparable to or greater than that observed in WT cells (**Figure 3B, iv and Figure 3C and 3D**). In contrast, the GTPase-defective ObgE(S173N) mutant showed substantially reduced rescue relative to WT ObgE. Thus, the ability of ObgE to compensate for the absence of Rbg1 depends on protein’s GTPase activity and an intact GTPase cycle. Although the absolute differences were modest, they were reproducible across independent experiments and statistically significant (**Figure 3D**). Together, these results demonstrate that bacterial ObgE can functionally substitute for yeast Rbg1 in supporting productive translation through a polyarginine-induced translational impediment. The ability of both proteins—but not their corresponding GTPase-defective variants—to promote downstream reporter synthesis supports conservation of a GTPase-dependent translation function across the bacterial and single-cell eukaryotic branches of the Obg family.

### 4. Bacterial ObgE GTPase promotes cell growth and mRNA translation through a polylysine sequence in human Drg1-deficient cells

Having found that bacterial ObgE can substitute for Rbg1 in yeast, we next asked whether this conserved activity extends to mammalian cells. Testing both systems was important for two reasons. First, complementation in yeast establishes functional conservation between bacterial ObgE and a eukaryotic Rbg protein in a genetically tractable organism. Second, complementation in human cells provides a more stringent cross-kingdom test and determines whether bacterial ObgE can operate within the substantially more complex mammalian translation environment.

First, we measured viable-cell metabolic activity in wild-type and Drg1-deficient HEK293T cells using the Alamar Blue assay. Whereas wild-type and Δrbg1 yeast cells displayed little difference in growth under standard conditions and only developed a detectable phenotype only in the presence of anisomycin, Drg1-deficient HEK293T cells exhibited a noticeable growth defect even in the absence of drug treatment (**Figure 4A**). The reduced alamarBlue signal in Δdrg1 cells was partially restored by expression of homologous human Drg1. Remarkably, expression of bacterial ObgE produced a comparable degree of rescue, indicating that ObgE can functionally substitute for human Drg1 despite their substantial evolutionary divergence. In contrast, the GTPase-deficient hDrg1(S78N) and ObgE(S173N) mutants failed to rescue the Δdrg1 phenotype. Thus, Drg1 makes a greater contribution to basal cellular fitness in HEK293T cells than Rbg1 does under standard yeast growth conditions, while the ability of bacterial ObgE to compensate for Drg1 loss supports a conserved, GTPase-dependent cellular function across bacteria, fungi, and mammals.

**Figure 4.**
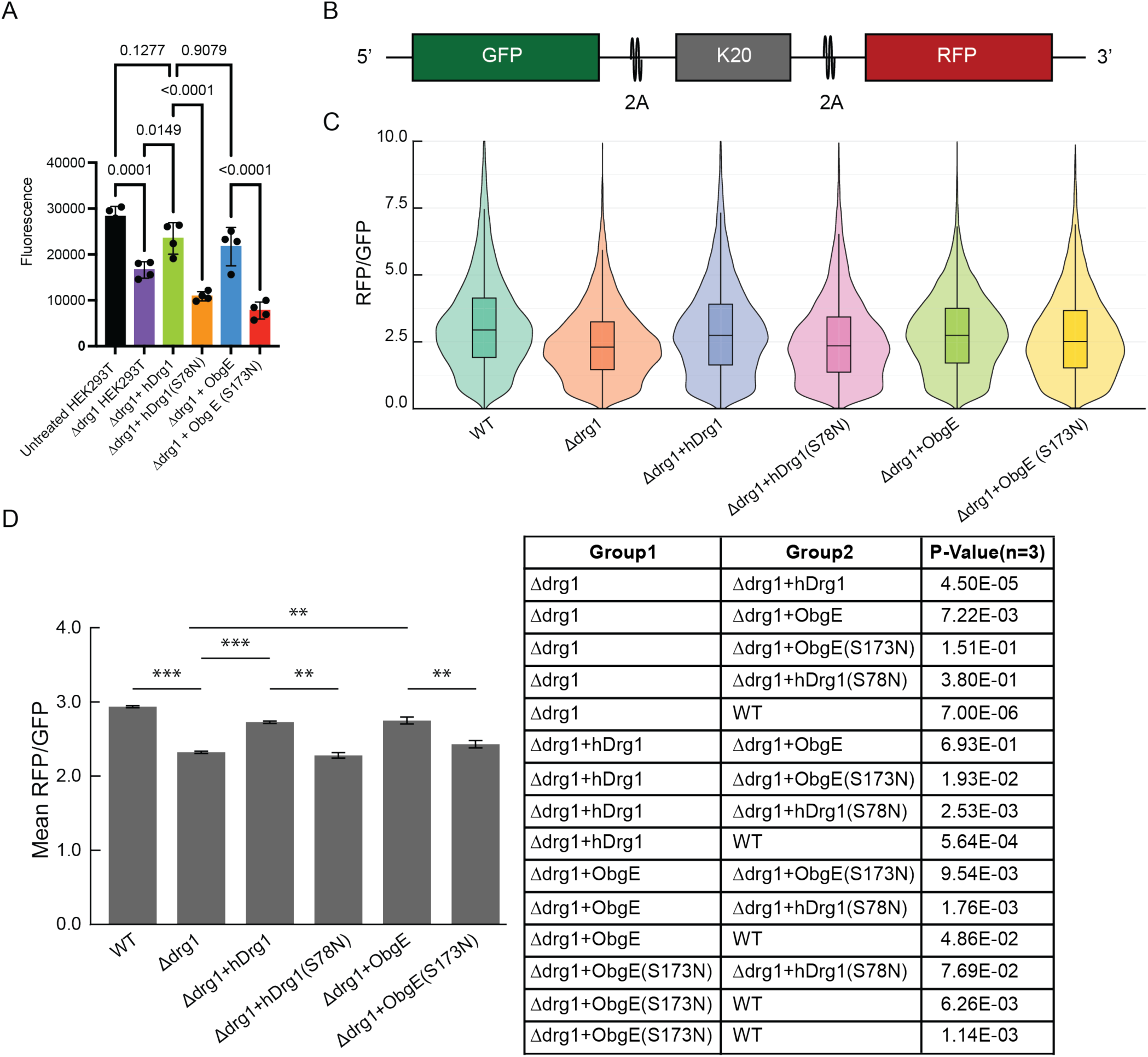
Bacterial ObgE GTPase promotes cell growth and mRNA translation through a polylysine sequence in human Drg1-deficient cells. **A. Bacterial ObgE restores metabolic growth activity in Drg1-deficient human cells.** HEK293T wild-type and Drg1-knockout (Δdrg1) cells were analyzed using the Alamar Blue assay, which measures cellular reduction of resazurin as a proxy for metabolically active cell number and growth. Δdrg1 cells were complemented with wild-type human hDrg1, GTPase-defective hDrg1(S78N), bacterial ObgE, or GTPase-defective Obg E (S173N). Drg1 deletion significantly reduced fluorescence relative to untreated wild-type HEK293T cells (*P* = 0.0001). Expression of wild-type human hDrg1 increased the signal in Δdrg1 cells (*P* = 0.0149), whereas mutant Drg1(S78N) failed to provide comparable rescue (*P* < 0.0001 versus wild-type DRG1). Expression of bacterial ObgE restored the signal, whereas ObgE (S173N) failed to rescue (*P* < 0.0001 versus wild-type ObgE). Individual points represent independent biological replicates; bars show the mean, and error bars indicate SD/SEM??. Statistical significance was determined using ???statistical test, with exact *P* values shown above the indicated comparisons. **(B-D) Bacterial ObgE GTPase promotes translation through a polylysine sequence in human Drg1-deficient cells** **B. Schematic of the dual-fluorescent reporter used to monitor ribosomal stalling in human cells.** The construct encodes GFP (eGFP) and RFP (mCherry) flanking a polylysine (K20) stalling sequence, separated by 2A sequences that allow independent release of each fluorescent protein. Ribosomal stalling at the stalling sequence reduces cellular RFP expression relative to GFP, enabling quantification via flow cytometry. **C. Bacterial ObgE rescues the translation defect caused by the loss of human Drg1.** Violin plots of single-cell RFP/GFP ratios from HEK293T cells co-transfected with the dual-fluorescent reporter construct and the indicated expression vectors. *Δdrg1* cells exhibit increased stalling, while plasmid-derived expression of WT hDrg1 or *E. coli* ObgE restores readthrough of the stalling sequence. GTPase-deficient mutants hDrg1 (S78N) and ObgE (S173N) fail to fully rescue readthrough. Each violin represents distributions of data from 23,000 individual cells from one representative replicate. **D. Stalling rescue across replicates.** The bar graph shows the mean of median RFP/GFP ratios from three replicates (23,000 cells per replicate). Translation past the K20 stalling sequence is statistically significantly reduced in *Δdrg1* cells relative to WT, and is statistically significantly restored by expression of WT hDrg1 or ObgE. Cells expressing GTPase-deficient mutants hDrg1(S78N) and ObgE(S173N) demonstrate statistically significantly diminished K20 translation readthrough. *=*P* < 0.05, **=*P* < 0.01, and ***=*P* < 0.001. *P*-values for all comparisons are shown in the table. *P*-values were calculated using a two-sided t-test on replicate-level medians.

Second, to extend the specific functional conservation from a single-cell eukaryotic organism to multicellular metazoans and to identify whether this GTPase shares a related translational function in human cells, we examined whether their bacterial homologs can also promote translation through a known impediment sequences by monitoring the degree of translation of a reporter in the *Δdrg1* HEK293T cells expressing Drg1 homologs from different species.

A similar dual-fluorescent reporter system, where EGFP (GFP) and mCherry (RFP) genes surround a poly(A) sequence (encoding poly-lysines, the K20 stalling sequence [**Figure 4B**]), allows quantification of the relative levels of translation through K20 within cell populations at a single-cell resolution (35). According to the experimental design, ribosomes that stall at the poly(A) sequences are unable to proceed to translate the subsequent RFP gene, thereby decreasing the cellular concentration of RFP relative to GFP. Due to the subsequent 2A sequence, GFP is released prior to K20-induced ribosomal stalling events or potential mRNA degradation by the downstream quality control, so the amount of cytoplasmic GFP reflects the degree of translation on the reporter before translating ribosomes reach the K20 sequence (**Figure 4C)**. As such, the ratio of RFP/GFP fluorescence intensity obtained in individual cells allows comparison of translation dynamics at such K20 impediment sequences between cell populations, as visualized through violin plots (**Figure 4D**). Translational machinery in *Δdrg1* HEK293T cell populations exhibits decreased translation past the poly(A) sequence compared to the WT cells expressing chromosomal Drg1, as indicated by a statistically significant ∼26% decrease in the median measured RFP/GFP ratio of Δ*drg1* cells versus WT (**Figure 4C and 4D**). This result suggests that human Drg1, similarly to its yeast homolog, Rbg1, promotes translation in the ribosome when ribosomes stall. To confirm that changes in RFP/GFP ratios reflected readthrough of the K20 stalling sequence rather than intrinsic differences in translation or fluorescence between cell types, K20 readthrough was compared to K0 controls in both WT and *Δdrg1* cells. The results showed that Drg1 knockout selectively worsened translational defects in the presence of the K20 stalling sequence (**Supplemental Figure 5**). We can not rule out a possible nucleic function of ObgE in eukaryotic cells, however, it is important to note that by normalizing the RFP signal to the GFP signal from the same reporter mRNA, we largely rule out changes in the fluorescence readout due to variations in the number of translation-competent ribosomes and reporter mRNAs between different cells.

While chromosomal knockout of human drg1 resulted in obvious ribosomal stalling on the poly(A) sequence, translation through the poly(A) stalling sequence was restored when *Δdrg1* cells were transfected with hDrg1– and bacterial ObgE-encoding vectors. As shown in **Figure 4C** and **4D**, the median RFP/GFP ratios of *Δdrg1* HEK293T cells increased by 17% and 18% upon expression of hDrg1 and E coli ObgE proteins, respectively. Median WT RFP/GFP ratios were approximately 8% and 7% higher than Drg1– and ObgE-supplemented *Δdrg1* HEK293T cells, respectively. These results suggest that the function of Drg to promote efficient translation is likely conserved across multiple domains of life.

To confirm that Drg’s GTPase-dependent activity is required to alleviate translational stalling, in addition to restored cell growth in the presence of anisomycin (**Figure 2A** line 4, and **1E** light blue and yellow lines), we mutated the conserved serine in the G1-motif of hDrg1 (S78N) and ObgE (S173N), to disrupt nucleotide binding thereby diminishing the GTPase activity in the protein. In both cases, the expression of hDrg1(S78N) and ObgE (S173N) decreased K20 translation relative to the expression of WT hDrg1 and ObgE proteins in *Δdrg1* HEK293T cells. Biological replicates of this set of experiments are included in **Supplemental Figure 6** to confirm the consistency of the observed RFP/GFP ratios in these cells. Thus, the bacterial protein can function within human cells, but its ability to restore downstream reporter synthesis depends on its GTPase activity.

Together with the yeast R12 reporter results, these findings demonstrate cross-kingdom functional interchangeability of Obg-type GTPases in two distinct eukaryotic systems and with two different polybasic translational impediments. In yeast, bacterial ObgE compensated for loss of Rbg1 during translation through consecutive arginine codons; in human cells, ObgE compensated for loss of Drg1 during translation through a polylysine sequence. The concordant results across organisms and reporter contexts support the conclusion that ObgE and Rbg1/Drg1 retain an evolutionarily conserved, GTPase-dependent activity that promotes productive translation through difficult coding sequences.

### 5. Obg-type GTPases promote puromycin incorporation in the stalled ribosome

It is striking that bacterial Obg supplementation can facilitate human ribosome translation through a poly(A) sequence. This finding is particularly unexpected because eukaryotic Rbg/Drg proteins function as heterodimeric GTPases on translating ribosomes through tight association with their partner proteins (3, 29), whereas bacterial Obg is a monomeric GTPase.

How, then, do these distinct GTPases facilitate efficient translation over an impediment sequence? Notably, bacterial Obg lacks the HTH, TGS, and S5D2L domains that are characteristic of eukaryotic Drg proteins, but possesses a unique N-terminal Obg-fold. The major structural feature shared between them is the conserved GTPase domain, which directly associates with the ribosomal GAC. Based on our structural analyses, we propose that Rbg/Drg GTPases protect physiologically paused ribosomes by, at least, preventing premature engagement of ribosome quality control factors. For example, the presence of Rbg1 on the ribosome sterically occludes binding of Gcn2—a stress-responsive kinase that phosphorylates Ser51 on eIF2α, triggering integrated stress response (36–38).

In addition to a blocking role as indicated by structural evidence, we wonder if direct binding of these GTPases to the ribosome promotes productive translation beyond the impediment/stall. Biochemical and structural evidence obtained from the characterization of 80S ribosomes bound with yeast Rbg1/Tma46 complexes led to the hypothesis that Rbg/Drg GTPases promote protein synthesis by stabilizing a paused or stalled translation apparatus in a translationally productive conformation. To test this hypothesis and further define the mechanism of Drg action, we examined puromycin reactivity of ribosomes stalled on a poly(A) sequence *in vitro* using a defined *E. coli* translation system that lacks ribosome biogenesis, quality control, protein release, and recycling factors. The system provides a simplified and tractable biochemical model to mimic ribosomal pause in a physiological context.

Since premature 3’ truncation of mRNA is an established means to stall elongating ribosomes reliably at defined loci (39), we used an mRNA construct lacking an in-frame stop codon, but containing either 0 or 20 consecutive AAA codons at the 3’ terminus to stall elongating ribosomes with either K0 or K20 tracks occupying their exit tunnels (**Figure 5A**).

**Figure 5.**
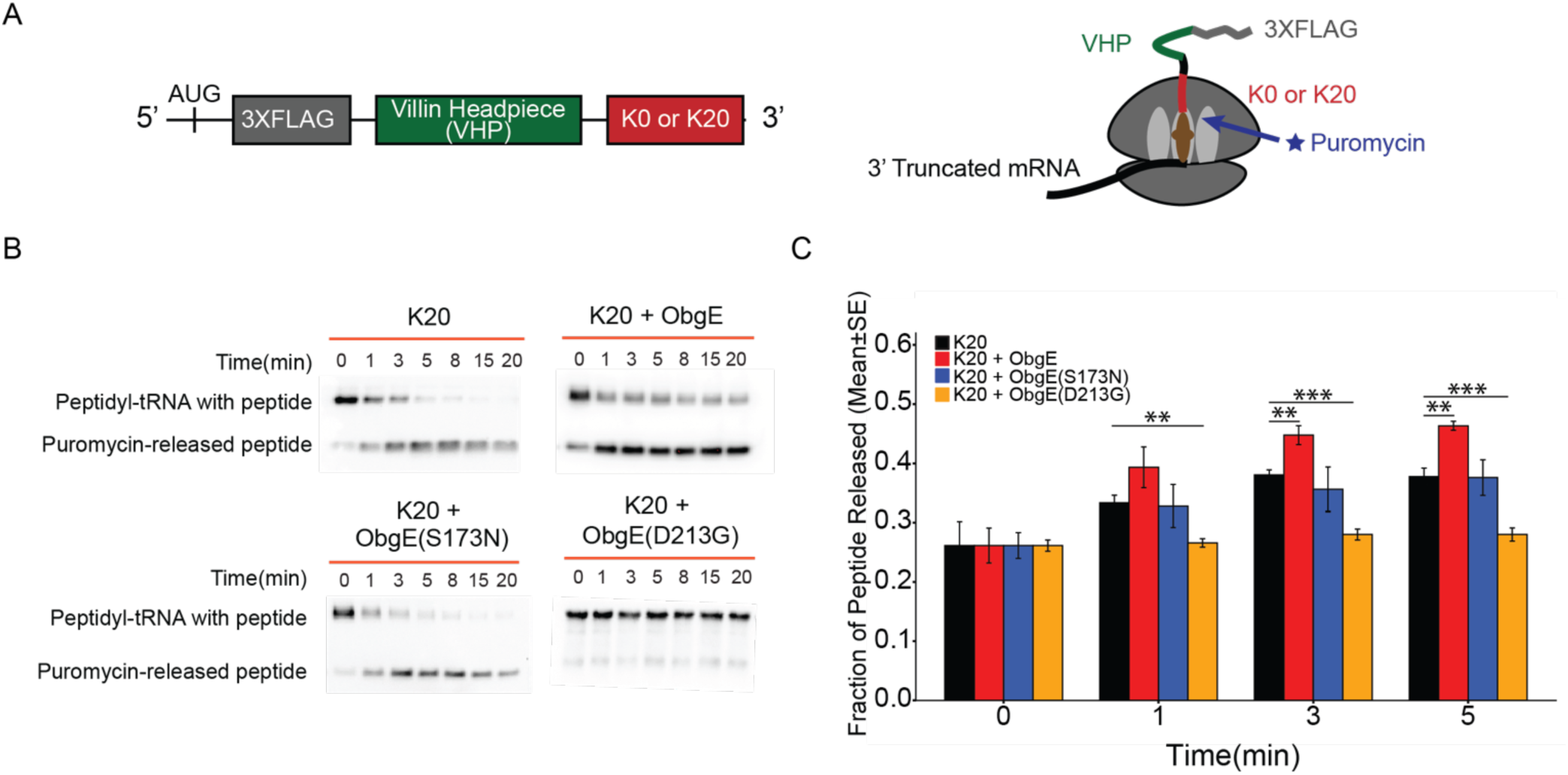
*E. coli* ObgE facilitates peptide bond formation in stalled ribosomes. **A. Schematic of the mRNA constructs used in the puromycin incorporation assay.** Ribosomes were stalled at the truncated 3’ terminus of mRNAs containing either 0 (K0) or 20 consecutive AAA codons (K20) in a defined reconstituted *in vitro E. coli* translation system devoid of release, recycling, and quality control factors. 2uM puromycin was added to all reactions to measure peptidyl transferase function in the ribosome by quantifying puromycin-conjugated nascent peptides. Either recombinant ObgE or an equal volume of buffer was added concurrently with puromycin to assay the ability of ObgE to facilitate peptide bond formation in the stalled ribosomes by the K20 sequences. **B** and **C. ObgE increases puromycin incorporation in K20-stalled ribosomes.** (B) Representative αFLAG immunoblots showing puromycin incorporation into nascent peptides. (C) Quantification of puromycin incorporation and release from at least three replicates of K20-stalled ribosomes. Enhanced peptidyl transferase activity is observed with wild-type ObgE incubation over 0-20 minute period (at 0,1,3,5,8,15 and 20 mins) post-puromycin addition. Within the same time-frame, ObgE (S173N) mutant, which is incapable of nucleotide-binding, shows no effect (blue line), and ObgE (D213G) mutant, which is incapable of GTPase-activity, inhibits puromycin-incorporation (orange line). Error bars represent the standard error of the mean.

It was shown that positively charged lysine stretches encoded by the poly(A) sequence induce conformational changes in the ribosome that disfavor peptidyl transfer (40, 41). To assess whether Obg/Rbg/Drg proteins alleviate translational stalling by promoting elongation-competent conformations in the ribosome, we monitored puromycin incorporation by stalled ribosomes. Since puromycin mimics the 3’ adenosine in the CCA-end of an aminoacylated tRNA charged with a modified tyrosine, it has been used as a direct measure of the geometric optimality of the peptidyl transferase center (PTC). As a CCA-structural mimic, puromycin can bind to the ribosomal A site. The reactive moiety of puromycin contains the requisite nucleophilic amine to attack the carbonyl carbon adjacent to the ester bond connecting the nascent peptide to the tRNA in the ribosomal P site in a manner identical to peptidyl transfer. Therefore, a comparison of the levels of puromycin incorporation into nascent chains within different ribosomal complexes will inform about relative differences in the capacities of their active sites (i.e., the PTC) to catalyze peptidyl transfer. As such, if the nascent chain conformation disfavors peptide bond formation due to a positively charged polylysine track in the exit tunnel, reduced puromycin reactivity is observed (40). In the presence of Obg/Rbg/Drg proteins, increased puromycin reactivity will demonstrate favorable conformational changes in the enzyme-substrate complex that are induced by this GTPase to compensate for suboptimal substrate orientation within a stalled ribosome.

Using an *E. coli*-derived *in vitro* translation system that contains *only* initiation and elongation factors, we assayed whether ObgE can target stalled ribosomes to promote puromycin incorporation ***at a 1:1 stoichiometry of ObgE and the stalled ribosomes***. Of note, puromycin was added concurrently with ObgE, while the ribosomes had already been stalled at the truncated 3’ mRNA termini, and the level of puromycin-conjugated peptides was measured over time. As expected, ObgE demonstrated a modest, yet consistent ability to facilitate puromycin incorporation in ribosomes that were translating the poly(A) sequence. As shown in **Supplemental Figure 7A**, comparing the translation of 3’truncated mRNAs lacking poly(A) encoded polylysine tracks (the K0 control) with mRNAs encoding 20 consecutive lysines (K20), the ability of K20-stalled bacterial ribosomes to catalyze puromycin incorporation was severely impaired. However, ribosomes translating the K20 sequence in the reaction mixtures containing *<u>equimolar</u> ObgE:70S* concentrations exhibited approximately 15-20 % increase in puromycin incorporation over the time course of 0-20 minutes when compared to otherwise identical reactions lacking ObgE (**Figure 5B**). In contrast, the mutant ObgE proteins, ObgE (S173N) mutant that is deficient in nucleotide binding (**Figure 5B and 5C)** and the ObgE (S173N-D213G) mutant that is deficient in both nucleotide-binding and GTP-hydrolysis (**Supplemental Figure 7B)**, fail to stimulate puromycin incorporation in the stalled ribosome. Remarkably, the ObgE (D213G) mutant, which is defective in hydrolyzing GTP but binds GTP, exhibits markedly reduced puromycin incorporation (**Figure 5B and 5C**, orange bar). This finding demonstrates that GTP hydrolysis is critical for puromycin incorporation. Biological replicates of the puromycin assay are reported in **Supplemental Figure 7C**. Since the amount of puromycin incorporated serves as a direct indicator of the level of peptide bond formation in the ribosome, this result supports a conserved molecular function of Obg-type GTPases to stabilize the stalled ribosome in a productive conformation ready for peptidyl transfer in the PTC in a GTPase-dependent manner. In line with this, the presence of human Drg1/Dfrp1 increases puromycin incorporation in the stalled ribosome using mammalian in vitro translation assays (16), suggesting a likely shared molecular mechanism for enhanced translation by this family of GTPase in eukaryotic cells

### 6. Rbg1-bound ribosomes preferentially associated with cellular mRNAs enriched for growth under our experimental condition

#### 6.1 A diverse set of endogenous cellular mRNAs associate with Rbg-bound translating ribosomes

The reporter and puromycin-incorporation experiments established that ObgE and Rbg1/Drg1 share a conserved capacity to promote productive protein synthesis. However, these experiments did not identify the endogenous messages on which Rbg1 normally acts. We therefore sought to define the physiological translation landscape associated with Rbg1.

Since Rbg1 binds to the ribosome, the presence of this protein increases mRNA stability and mRNA translation (3), we proceeded to define the mRNA population physically associated with Rbg1-containing ribosome complexes by isolating Rbg1-containing polysomes and sequencing their associated mRNAs. In this experiment, we isolated ribosome fractions, immunoprecipitated Rbg1-associated complexes, and performed mRNA-seq on the recovered RNAs. (**Figure 6A**). Specifically, yeast cells with chromosomally FLAG-tagged Rbg1 were grown at 30°C until they reached an OD_600_ of 0.6, then were treated with 100 µg/ml of cycloheximide for 5 minutes before harvesting. Polysome profiles were performed using 10%– 50% sucrose gradients. Fractions for 80S and polysomes were pooled, and the ribosomes were pelleted down by ultracentrifugation. Resuspended ribosomes were first incubated with magnetic FLAG beads, followed by extensive washing using polysome buffer three times. RNAs were extracted and subjected to mRNA-seq (42).

**Figure 6.**
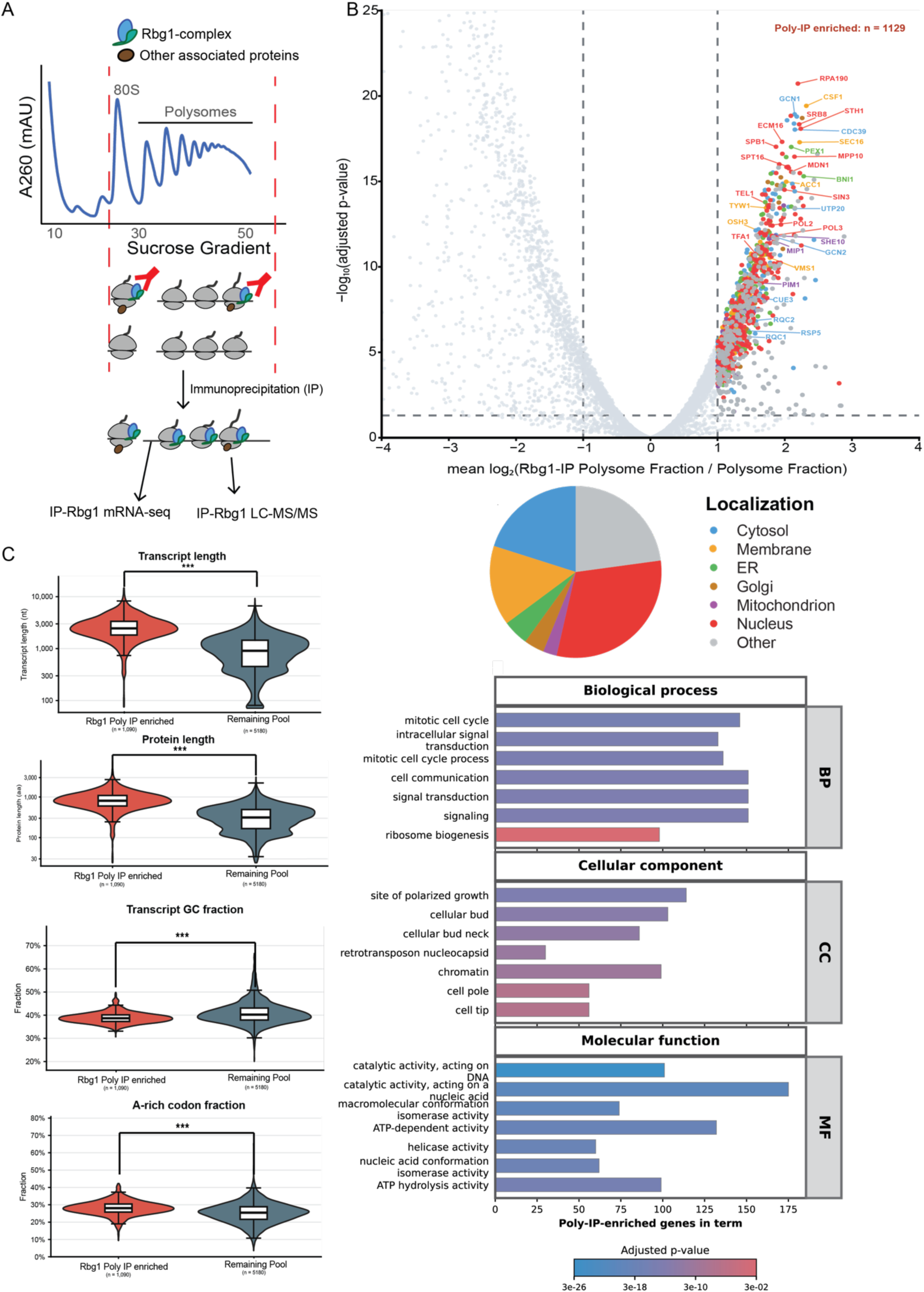
Identification of mRNAs associated with Rbg1-80S translation complex by IP-mRNA sequencing. **A. Schematic of Rbg1-IP mRNA-Seq and LC-MS/MS.** Rbg1-IP mRNA-Seq used to define the mRNAs associated with the Rbg1-bound ribosomes. 80S and polysome fractions were collected from sucrose gradients prepared from lysates of yeast expressing TAP-tagged Rbg1, and subjected to immunoprecipitation with IgG beads. Bead-bound RNAs were extracted, poly(A)-containing mRNAs were captured by oligo-dT, and subjected to mRNA-Seq. Similarly, IgG bead-bound proteins were digested, subjected and analyzed by LC-MS/MS. **B. Identification and functional classification of Rbg1 Poly-IP-enriched transcripts.** The volcano plot compares Rbg1 Poly-IP RNA recovery with matched polysome input using DESeq2-normalized protein-coding gene counts. Enrichment was defined as log2 fold change greater than 1 and Benjamini–Hochberg-adjusted p<0.05, identifying 1,129 transcripts. Selected genes involved in ribosome biogenesis, transcription, cell-cycle progression, membrane trafficking, and translation quality control are labeled. Points indicate UniProt localization; nonsignificant genes are light gray, and dashed lines mark statistical thresholds. The pie chart summarizes enriched-protein localizations. Gene Ontology analysis identified mitotic cell-cycle regulation, intracellular signaling, polarized growth, ribosome biogenesis, cellular-bud and chromatin components, and nucleic-acid-directed catalytic, ATP-dependent, helicase, and conformational-isomerase activities. Bar length indicates gene count and color indicates adjusted p value. **C. Sequence and protein features.** Poly-IP-enriched transcripts were longer than the remaining tested pool (median 2,450 versus 909 nucleotides) and encoded longer proteins (809 versus 312 amino acids). They also showed lower coding-sequence GC content and higher A-rich-codon fractions, defined as codons containing at least two adenines. Violin plots show distributions, interquartile ranges, and medians. Two-sided Wilcoxon rank-sum tests with Benjamini–Hochberg correction assessed significance; *** denotes adjusted p<0.001. Together, these data define a distinct Rbg1-associated transcript class enriched for long, A-rich messages supporting ribosome production and proliferative growth programs.

After read alignment and gene-level quantification, we verified data quality. Polysome fraction mRNA-seq and Rbg1-immunoprecipitated polysome fraction mRNA-seq data were reproducible across biological replicates, with correlations of 95.1% and 96.8%, respectively (**Supplemental Figure 8A and 8B**; **Dataset S1**), Transcript abundance in the Rbg1 immunoprecipitate was compared with that in the matched total polysome fraction, thereby distinguishing preferential Rbg1 association from overall transcript representation in translating ribosomes (**Supplemental Figure 8C, Dataset S2**). This analysis identified 1,129 transcripts significantly enriched in the Rbg1 polysome immunoprecipitated (**Figure 6B**). Enriched transcripts were defined using thresholds of log₂ fold enrichment greater than 1 and adjusted *P*-value below 0.05. The broad distribution of enrichment values indicated that Rbg1 association was not uniform across polysome-associated mRNAs but instead varied substantially among endogenous transcripts. Several of the most strongly enriched messages encoded proteins involved in transcription, chromatin regulation, ribosome biogenesis, signaling, and intracellular organization, including RPA190, GCN1, CDC39, SRB8, STH1, SEC16, SPB1, ECM16, and UTP20.

The proteins encoded by Rbg1/80S-associated transcripts were predicted to localize to multiple cellular compartments (**Figure 6B**, pie chart). Nuclear proteins constituted the largest annotated group, followed by proteins assigned to the cytosol and cellular membranes. Smaller subsets were associated with the endoplasmic reticulum, Golgi apparatus, and mitochondria. Thus, Rbg1 association is not restricted to translation products destined for a single organelle. Instead, Rbg1-containing polysomes synthesize proteins with diverse cellular destinations, consistent with Rbg1 acting on cytosolic ribosomes according to the demands of translation given the growth condition rather than the ultimate localization of the protein product.

Gene Ontology analysis further revealed that the Rbg1-associated transcript population was enriched for biological processes connected to cell growth and proliferation, including the mitotic cell cycle, signal transduction, cell communication, and ribosome biogenesis (**Figure 6B**). Cellular-component annotations included sites of polarized growth, the cellular bud and bud neck, chromatin, and the cell pole or tip, suggesting that Rbg1-associated translation contributes to processes requiring coordinated cell-cycle progression and spatial organization. Molecular-function categories were dominated by ATP-dependent and nucleic-acid-associated activities, including catalytic activity acting on DNA or other nucleic acids, helicase activity, ATP hydrolysis, and macromolecular or nucleic-acid conformation isomerization.

#### 6.2 Rbg1-bound ribosome preferentially associates with long, A-rich mRNA transcripts when the cell is undergoing fast growth in rich media

Having identified a selective set of endogenous mRNAs associated with Rbg1-containing polysomes, we next asked whether these transcripts share sequence or protein-coding features that might explain their preferential association with Rbg1/80S. We compared the 1,090 Rbg1-enriched mRNA transcripts versus the mRNA detected in the translation pool (**Figure 6C**). Under our current growth condition, Rbg1/80S-associated transcripts were longer than those in the unenriched pool. Their median transcript length was 2,454 nucleotides, compared with 909 nucleotides for other transcripts, representing a 170.0% increase. This difference was reflected in the encoded proteins: proteins produced from Rbg1-associated transcripts had a median length of 809 amino acids, compared with 312 amino acids for the remaining population (Cliff’s δ = 0.757, adjusted *P* < 1 × 10^−300^) (**Figure 6C**).. Thus, Rbg1 preferentially associates with polysomes translating long open reading frames.

Furthermore, the secondary structure of the mRNA does not seem to play a critical role, as suggested by the a similar GC content of enriched Rbg1/80S-associated mRNA (38.67%) versus the remaining pool (40.24%). In contrast, Rbg1/80S-associated coding sequences contained a higher fraction of A-rich codons, which were defined as codons containing at least two adenosines (28.08% versus 25.48%; Cliff’s δ = 0.318, adjusted *P* = 4.43 × 10^−61^). These results suggest that Rbg1-bound ribosomes are associated not only with mRNA transcript length but also with a distinct nucleotide composition with A-rich codons.

### 7. A diverse range of protein peptides associate with Rbg-bound translating ribosomes

The selective transcript association suggested that Rbg1 may operate within specific co-translational environments under the standard growth conditions of the study. We therefore asked whether the protein composition of Rbg1/80S-associated complexes reflected the cellular destinations of the encoded proteins and the demands imposed by their synthesis. To gain insight into the repertoire of nascent peptides whose translation is facilitated by Rbg/Drg association, and to identify additional proteins that interact with Rbg/Drg-bound translating ribosomes, we performed immunoprecipitation–mass spectrometry (IP–MS) analysis of yeast Rbg1-bound translation complexes, to gain a complementary protein-level view of these complexes. In this experiment, we compared the relative abundances of proteins detected in Rbg1-enriched ribosomal fractions to those in total cellular ribosomes (unenriched samples) using LC–MS/MS. Cellular ribosomes were purified from *S. cerevisiae* strains harboring a chromosomally integrated C-terminal 3×FLAG tag on Rbg1 via sucrose cushion sedimentation. Rbg1-associated ribosomes were subsequently enriched from these preparations by αFLAG immunoprecipitation (**Supplemental Figure 9A** and **Figure 7A**). Experimental controls were carried out (**Supplemental Figure 9B)**. It’s worth mentioning that we also carried out salt wash with high KCl concentrations to confirm that the Rbg1-IP pulls down ribosomes, translation factors and nascent chain complexes (**Supplemental Figure 9C).** A similar pulldown method was used to determine the cryoEM structure of 80S ribosome bound to Rbg1/Tma46 (3)

**Figure 7.**
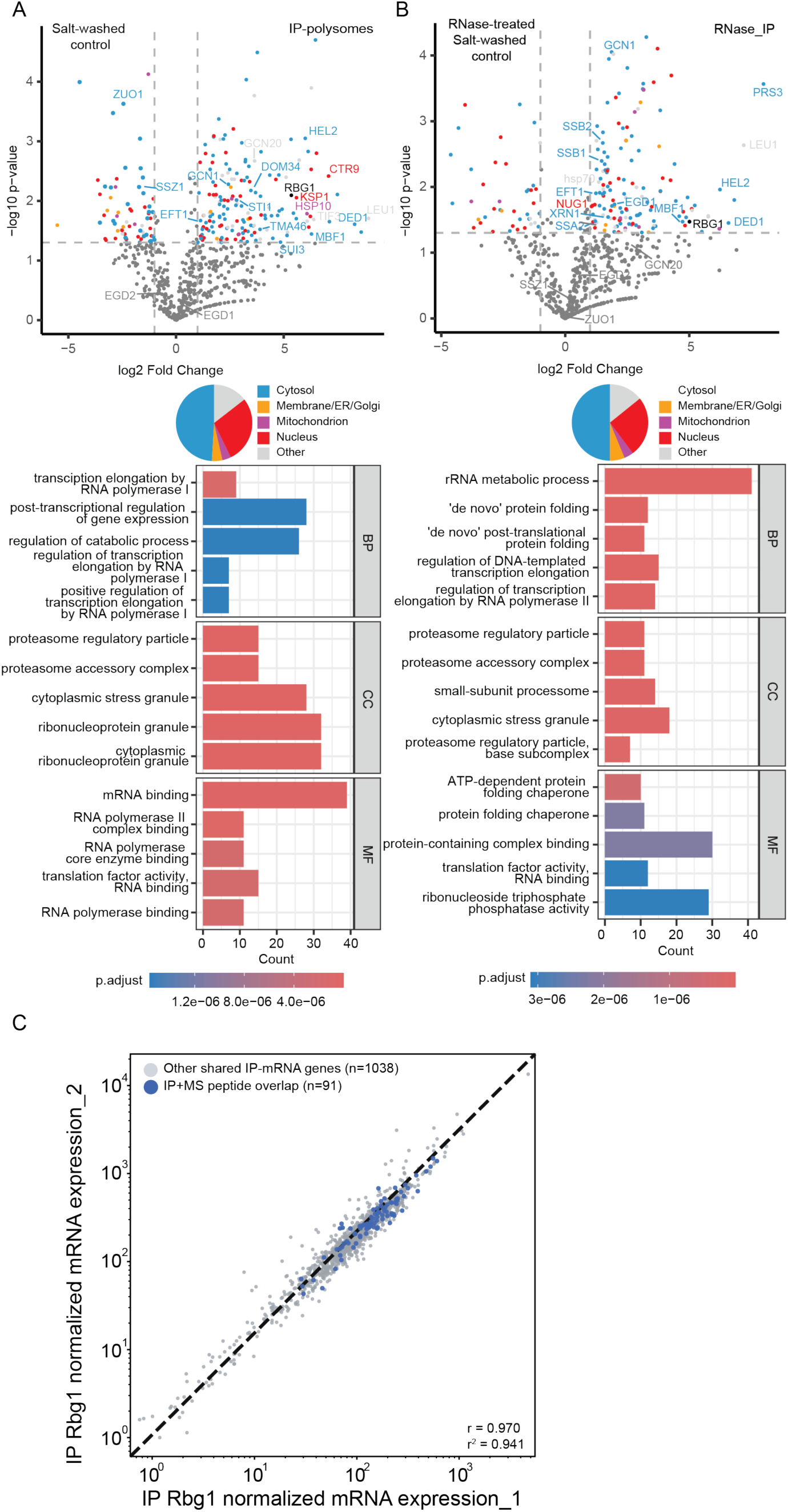
Identification of peptides associated with Rbg1-80S translation complex by IP-LC-MS/MS. **A** and **B. Enriched and de-enriched peptides in the Rbg1-IP samples relative to proper salt-washed control of cellular 80S ribosomes.** Volcano plot showing significantly enriched or de-enriched peptides, as determined by log_2_ fold change >1 or <-1, respectively with adjusted *p*-value <0.05 (Benjamini-Hochberg). Volcano plots display proteins quantified in salt-washed control versus IP-polysomes (**A**) and RNase-treated and salt-washed control versus RNase-treated IP (**B**), plotted as log₂ fold change against – log₁₀ *p*-value. Statistical thresholds (vertical and horizontal dashed lines) were used to define significantly enriched proteins. Highlighted proteins are color-coded according to their annotated subcellular localizations, and representative enriched proteins are labeled. Subcellular localization information from Uniprot was categorized into cytosol, membrane/ER/Golgi, mitochondrion, nucleus, and other compartments. GO enrichment analyses were performed on significantly enriched proteins only and are shown for Biological Process (BP), Cellular Component (CC), and Molecular Function (MF). Bars represent the number of proteins associated with each enriched GO term, and bar colors indicate adjusted p-values. **C: Correlation plot of Rbg1-IP mRNA enriched dataset and overlap with the IP-MS peptide dataset.** Normalized Rbg1-IP-mRNA replicate expression values are plotted on a log scale for all IP-mRNA-enriched genes (n = 1129). Genes shared between the IP-MS and IP-mRNA datasets (n = 91) are highlighted in blue. All remaining IP-mRNA genes (n = 1038) are shown in gray. The dashed diagonal represents the line of identity between replicates. Representative genes within the 91-gene overlap include SPT5, PAF1, CTR9, NOP14, RRP9, GCN1, RPN2, SEC16, and STE20. These genes are consistent with enrichment for factors involved in transcription, ribosome biogenesis, translation stress, protein turnover, vesicle trafficking, and cell signaling. Pearson correlation coefficient for all IP-mRNA genes: r = 0.970 (r² = 0.941).

**Figure 8.**
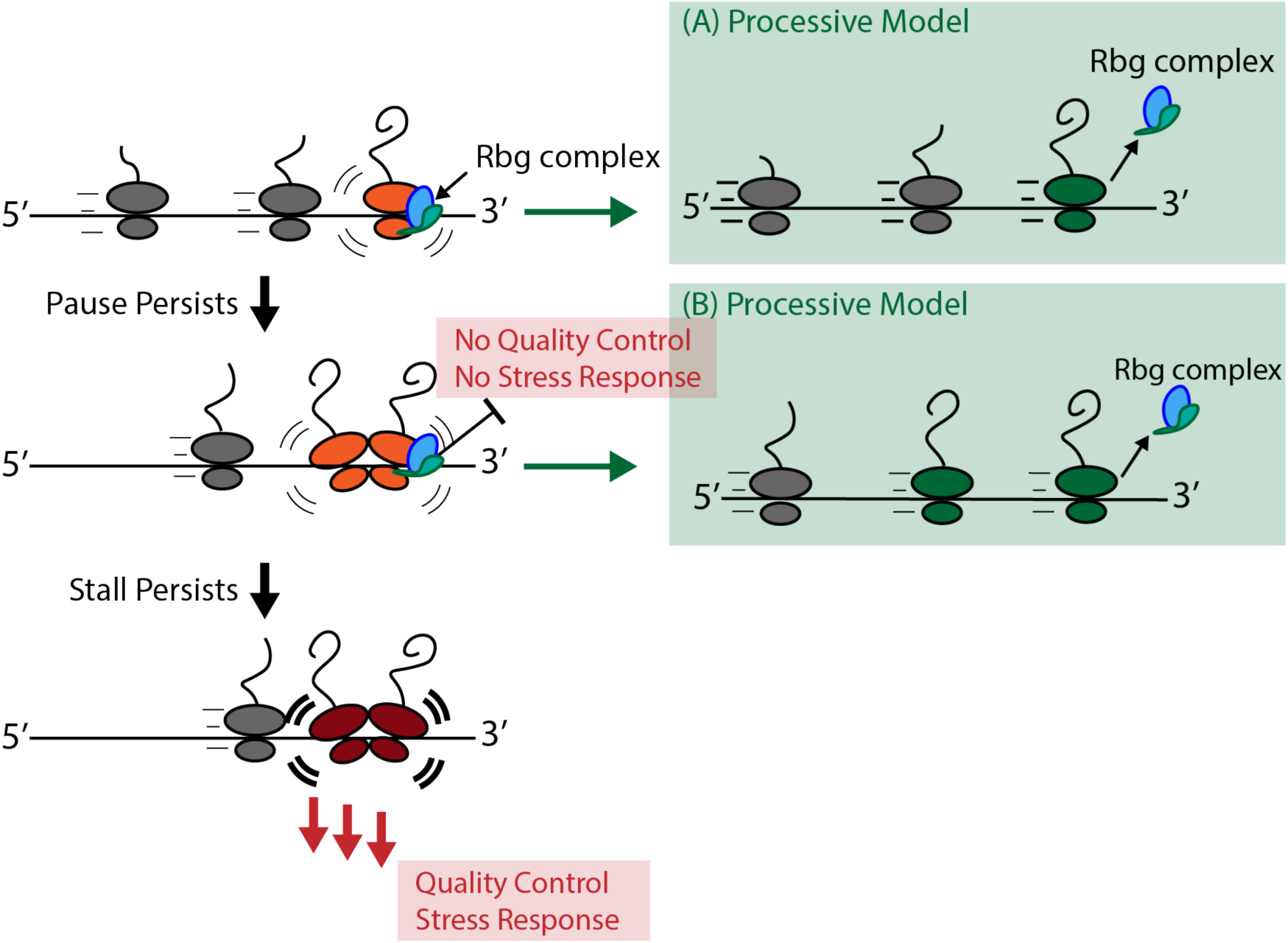
Function of Rbg/Drg GTPase in protein synthesis. Rbg/Drg GTPases bind to slowed or stalled translating ribosomes and promote translation, rather than downstream quality control and stress response pathways, thereby providing the ribosome an opportunity to overcome the stall before mRNA translation is prematurely and unproductively terminated. Rbg/Drg complexes target the slowed ribosomes, or collided disome, providing the problematic ribosome with more time to ‘decide’ whether to commit to quality control, all the while promoting the stalled ribosome or leading ribosome in the collided pair to continue translating (A and B, processive model). Rbg/Drg proteins can attenuate the ISR pathway by competing with Gcn2 for binding to the ribosome in a productive way (i.e., placeholder model). Finally, we cannot rule out the possibility that Rbg/Drg proteins may, at some point in this pathway, target the problematic ribosomes for splitting in a manner that preserves the integrity of the involved mRNA, although none of the current data support this model.

Relative peptide abundances between Rbg1-enriched and unenriched samples were estimated using emPAI scores calculated by Mascot, followed by normalization across samples. The average log₂-transformed amount for each protein was used to calculate log₂-fold changes, indicating enrichment or depletion in the Rbg1-bound translation complexes relative to the total polysomes (**Figure 7A**).

Moreover, to obtain a more accurate control that accounts for the potential loss of loosely ribosome-associated ribosomal proteins during the IP washing procedure, *<u>unenriched cellular polysomes</u> <u>were subjected to the same wash conditions</u>* as the IP samples. Specifically, sucrose-cushioned ribosomal pellets were resuspended in IP wash buffer containing 20mM Tris-HCl, pH 7.5; 150mM KCl; 5mM MgCl_2_; 2μM ZnCl_2,_ and incubated with end-over-end rotation for 20 minutes at 4 °C. The washed samples were then re-pelleted through a 1M sucrose cushion and resuspended in the same buffer used for IP-enriched fractions. This parallel treatment provided a rigorously matched control for evaluating the specificity of Rbg1-associated ribosome enrichment.

We identified 269 proteins that were significantly enriched in the Rbg1-bound ribosomal complexes relative to unenriched ribosomal fractions (*p*-value **<** 0.05; log₂ fold change ≥1). These enriched proteins represent a diverse spectrum of cellular functions (**Dataset S3**). Because MS method used here is only semiquantitative, we focused on proteins detected in both Rbg1-enriched and unenriched ribosomal fractions to ensure a more reliable comparison. Proteins detected exclusively in the Rbg1-enriched samples are included in **Dataset S3** for reference.

Among the proteins identified in both datasets, the statistically significant subset with the larger than 2-fold changes, 29% and 4% of these proteins correspond to factors with annotated nuclear or mitochondrial localization (**Figure 7A**), suggesting extensive crosstalk between cytoplasmic translation and gene expression pathways in these organelles. The enriched nuclear proteins span multiple categories, including nucleolar ribosome biogenesis, transcription, pre-mRNA splicing, cell cycle, and chromatin proteins. Nascent chain–associated proteins such as Zuo1 (a J-domain protein) and Ssz1 (a Hsp70-like cofactor) (43, 44) appeared in the de-enriched IP-polysome pool. Gene ontology (GO) analysis of proteins identified this way is shown (**Figure 7A**). Given that the yeast cells were harvested during exponential growth in nutrient-rich medium, it is perhaps unsurprising that nuclear and mitochondrial proteins involved were among the strongly represented. The enrichment of these proteins specifically in Rbg1-bound translation complexes suggests that Rbg1 plays a role in facilitating translation of organelle-targeted proteins, potentially coordinating cytoplasmic translation with organelle function and cell growth.

To determine whether the interactions between Rbg-associated ribosomes and regulatory factors are mediated by RNA or depend on the mRNA being translated, we treated the ribosomal fraction with RNase I prior to Rbg1-immunoprecipitation, followed by LC–MS/MS analysis (**Supplemental Figure 9A**). This approach not only eliminates potential RNA-mediated interactions but also enhances the representation of nascent-chain–associated peptides and co-translational factors that interact directly with the ribosome or emerging polypeptides.

In RNase I digested samples, we detected a broad set of translation factors and co-chaperones that associate with nascent-chain folding and targeting (**Figure 7B** and **Dataset S4**). For example, ribosome-associated chaperones Hsp70 family chaperones, Ssb1 and Ssb2, which play essential roles in co-translational protein folding, were significantly enriched. Sti1, a cytosolic co-chaperone linking Hsp70 and Hsp90, was consistently identified in the IP-enriched RNase-treated samples.

Importantly, in our mass spectrometry analysis of Rbg1-IP ribosomes, we detected Mbf1 (the yeast homologs of mammalian EDF1), Hel2 (the yeast homologs of mammalian ZNF598), Gcn1 among the co-purified proteins in both RNase-treated and non-RNase-treated samples. Gcn20 was also detected in all samples, although it does not meet our *p*-value cutoff after the RNase 1 treatment (log_2_ fold change = 2.497; *p*-value = 0.0734). These factors are well-known components of the ribosome collision–sensing (45–47) and translational stress-response machinery (36–38), and their presence in Rbg-associated ribosomal fractions provides key insight into Rbg’s functional context.

The presence of Mbf1, Hel2, Gcn1, among the co-purified proteins in the RNase I-treated samples suggests that these factors remain associated via ribosome-proximal or protein–protein interactions rather than through mRNA-mediated interaction, supporting a direct functional link between Drg-bound ribosomes and stress response/quality-control machineries.

The co-enrichment of these factors with Rbg suggests that *Rbg GTPases associate with ribosomes at a regulatory checkpoint where translational pausing, collision sensing, and stress signaling intersect.* This finding supports our model that Rbg/Drg proteins act upstream of RQC and ISR components, stabilizing physiologically paused ribosomes to prevent premature recruitment of stress-response machinery. In this view, Rbg/Drg binding marks a productive paused state, while Mbf1–Hel2 and Gcn1– Gcn20 represent coupled surveillance and stress response factors that are poised to engage further (such as recruiting Gcn2) if pausing persists. Of note, the protein peptides that were detected by IP-MS belong to a subset of the products that are translated from mRNAs associated with Rbg1-bound ribosomes (**Figure 7C**).

## Discussion

### Obg and Rbg/Drg represent ancient, functionally related branches of the Obg-like superfamily GTPases

Our findings support an evolutionarily conserved role for Obg/Rbg/Drg GTPases in cellular translation. Phylogenetic analysis resolved bacterial Obg and archaeal/eukaryotic Rbg/Drg as distinct, deeply diverged branches within the broader Obg-like GTPase superfamily. Archaeal organisms generally contain a single Rbg-like protein, whereas eukaryotic sequences separate into the Rbg1/Drg1 and Rbg2/Drg2 lineages, consistent with an early duplication preceding the diversification of major eukaryotic groups. The widespread retention of these proteins implies sustained selective pressure to preserve their cellular functions.

The phylogeny suggests Obg and Rbg/Drg likely arose through early divergence and specialization of an ancestral Obg-family GTPase. Their distinct domain architectures and evolutionary histories indicate substantial functional adaptation. Nevertheless, their common ancestry, ribosome association, and functional interchangeability (this study) suggest that they retain an ancestral biochemical activity related to protein synthesis. This distinction is important for understanding the extent of conservation, as sequence divergence does not necessarily indicate complete divergence of biochemical function. *Instead, the Obg-family scaffold may have preserved a core ability to sense and/or modify ribosomal states while acquiring lineage-specific domains, partners, and regulatory mechanisms. Our results suggest that this core activity has been maintained from bacterial Obg to eukaryotic Rbg/Drg*.

### Cross-species complementation reveals a conserved translation function

The ability of bacterial ObgE to substitute for Rbg1/Drg1 provides direct functional evidence that the relationship between these GTPases extends beyond sequence or structural similarity. In yeast, deletion of Rbg1 modestly but statistically significantly reduced downstream protein synthesis across a sequence encoding 12 consecutive arginines. Expression of either yeast Rbg1 or bacterial ObgE restored reporter output, whereas GTPase-defective variants failed to provide comparable rescue. Similarly, loss of Drg1 in human cells reduced translation through a sequence encoding 20 consecutive lysines, and this defect was rescued by either human Drg1 or bacterial ObgE, but not their GTPase-deficient counterparts.

Using both yeast and human cells provides complementary evidence. The yeast system establishes functional substitution in a genetically tractable single-cell eukaryote, whereas rescue in human cells provides a more stringent test of whether bacterial ObgE can operate within a divergent mammalian translation environment. The two reporters also contain different polybasic sequences, arguing that complementation is not restricted to one particular amino acid or codon sequence.

The magnitudes of the reporter effects were modest, but this is not unexpected for factors that facilitate translation under specific conditions rather than serving as essential components of every elongation/peptide-bond formation cycle. Rbg1/Drg1 may act on only a subset of ribosomes or during transient kinetic states. A population-level reporter would therefore integrate strongly affected ribosomes with many translation events that proceed independently of Rbg1/Drg1 (for example, **Figure 3**). The reproducibility, statistical significance, rescue by the wild-type proteins, and reduced activity of the GTPase-defective variants together support the biological relevance of these effects.

These reporters measure successful synthesis of a downstream protein and therefore provide evidence for productive translation beyond the polybasic sequence. They do not, by themselves, determine whether Rbg1/Drg1 accelerates decoding, tRNA accommodation, peptide-bond formation, translocation, or recovery from a paused state. Nevertheless, the cross-species rescue demonstrates that ObgE and Rbg1/Drg1 share a GTPase-dependent activity that increases the probability that translation will continue productively through difficult coding sequences in the mRNA.

### Obg GTPases stabilize the paused ribosome in a competent state poised for translation

Spatially and temporally regulated ribosomal pauses require active protective mechanisms that prevent premature engagement of quality control or stress response pathways. We propose that Drg-type GTPases serve this protective role inherent in the physiological context of the cell, where they bind and stabilize the translating ribosome in a competent state, ensuring that translation resumes efficiently once the ribosomal pause is over.

To experimentally evaluate this model, we sought to determine whether Obg-type GTPases actively stabilize paused ribosomes in a translationally competent conformation. Physiological ribosomal pauses occur naturally during processes such as signal-sequence recognition, membrane targeting, or organelle import, where the elongating ribosome must transiently slow translation to coordinate with downstream pathways. To mimic this controlled pausing in a simplified and well-defined biochemical context, we reconstituted ribosome stalling on poly(A) sequences *in vitro <u>using a purified E. coli</u> <u>translation system that is devoid of ribosome biogenesis, quality-control, and recycling factors.</u>*We then assessed the effect of Obg GTPases on nascent-chain reactivity with puromycin, providing a direct measure of whether GTPase binding promotes a translationally productive state beyond the stall. The degree of puromycin-incorporated into the nascent chain serves as a biochemical indicator of whether the peptidyl transferase center is properly aligned by the GTPase and whethethe ribosome is poised for elongation further. Using this method, we observe an increase in puromycin reactivity in the presence of ObgE at an equal molar ratio to the ribosome, suggesting that these GTPases stabilize the stalled ribosome in a conformation competent for peptide bond formation—an essential prerequisite for continued translation. Of note, the degree of puromycin incorporation at a 1:2 stoichiometry of the programmed ribosome and ObgE, as well as at higher concentrations of puromycin (e.g., 500 mM and 1 mM), was also measured, and a similarly increased level of incorporated puromycin was obtained.

Furthermore, the puromycin-incorporation experiments provide a biochemical connection between cross-species complementation and ribosomal catalysis. Puromycin enters the ribosomal A site and reacts the nascent polypeptide from the P-site peptidyl-tRNA. Its incorporation therefore requires an accessible and sufficiently reactive peptidyl-transferase center. Enhanced puromycin incorporation in the presence of ObgE indicates that ObgE increases the capacity of stalled ribosome–nascent-chain complexes to perform peptidyl transfer.

This activity offers a mechanistic explanation for the reporter rescue. Ribosomes encountering polybasic sequences can enter slowly reacting or unproductive conformational states. By binding such ribosomes, ObgE, in a GTPase-dependent manner, may stabilize or restore a conformation that supports accommodation of an A-site substrate and reaction with the P-site peptidyl-tRNA. Rbg1/Drg1 may perform an analogous function in eukaryotic cells, thereby protecting (as well as maybe even extending) the productive lifetime of a paused ribosome and allowing elongation to resume before the complex is committed to translational arrest or quality-control pathways.

Importantly, puromycin reactivity should not be equated with completion of the entire elongation cycle. Increased incorporation could reflect improved A-site accessibility, altered positioning of peptidyl-tRNA, enhanced peptidyl-transferase-center reactivity, or redistribution among ribosomal conformations. Additional kinetic and structural experiments will be required to distinguish these possibilities. Nonetheless, the puromycin results place the conserved activity of ObgE close to the catalytic center of the ribosome and support a role in promoting nascent-chain synthesis/peptide-bond formation rather than simply changing reporter expression indirectly.

While our *in vitro* assay is tested on an elongating ribosome, the conclusion obtained that Obg/Rbg/Drg-GTPase stabilizes the paused ribosome in a competent conformation can be extended to other phases in translation, including initiation and termination/recycling. Our IP-MS detected many initiation and termination factors, and our earlier genome-wide sequencing results demonstrated that this GTPase indeed functions in translation beyond the elongation phase (3).

### Rbg1 GTPase at the interface of mRNA translation, nascent-chain maturation, and proteostasis

Although the R12 and K20 reporters define specific translation challenges, endogenous Rbg1 function is unlikely to be limited to uninterrupted polybasic sequences. Rbg1 IP–mRNA-seq revealed that Rbg1 association is selective across the yeast translatome. Rbg1-containing polysomes were preferentially associated with long transcripts displaying an A-rich coding composition when the cell is undergoing fast growth in rich media. Gene Ontology analysis further connected these transcripts to ribosome biogenesis, nucleic-acid metabolism, signaling, cell-cycle progression, and polarized growth.

Additionally, Rbg1 IP-MS complements the transcriptomic analysis by defining the protein environment surrounding Rbg1-associated translation complexes. Under our experimental conditions, Rbg1-bound ribosomes were enriched with translation factors, RNA-binding proteins, ribosome-surveillance factors, chaperones, and proteasome-associated machinery. These proteins included factors connected to translational regulation and surveillance, such as Gcn1, Gcn20, Hel2, Dom34, Mbf1, and Ded1, as well as the Rbg1 partner Tma46. For example, Mbf1 and Hel2, key components of the ribosome collision–sensing and quality-control pathway (45–47), as well as Gcn1 and Gcn20, which initiate the integrated stress response when ribosomes stall (36–38). Importantly, enrichment of these factors (Mbf1, Hel2 and Gcn1, as well as Gcn20) in RNase I–treated Rbg1–IP samples suggests that their association with the Rbg1-bound ribosomes is likely to be *<u>RNA-independent</u>*, reflecting direct or ribosome-proximal interactions rather than nonspecific co-sedimentation through shared mRNAs. The detection of these factors, and the overall composition of enriched proteins in the Rbg1-immunoprecipitated samples, suggest that Rbg1 interacts with ribosomes positioned at regulatory or paused states rather than with constitutively elongating complexes. This result Rbg1 within a translation network that monitors ribosome activity and coordinates it with RNA metabolism and protein quality control.

RNase treatment provided a further important distinction. Disruption of RNA– and ribosome-mediated connections altered the Rbg1-associated proteome but preserved associations with several translation and protein-folding factors, including the ribosome-associated Hsp70 chaperones Ssb1 and Ssb2 and other chaperone-related proteins. The RNase-resistant population was enriched for *de novo* protein folding, chaperone activity, protein-complex binding, rRNA metabolism, and translation-factor activity. These findings suggest that at least part of the Rbg1 interaction network is maintained through protein-mediated or stable complex-level associations rather than passive co-occupancy on the same mRNA. The convergence between IP–mRNA-seq and IP-MS strengthens this interpretation. Hundreds of proteins recovered by IP-MS were encoded by transcripts reproducibly detected in the Rbg1 IP–mRNA dataset (**Figure 7C**). This overlap does not necessarily establish that every recovered protein represents a nascent chain being synthesized at the moment of immunoprecipitation; some may be stable interactors, chaperones, regulatory proteins, or mature protein products. Nevertheless, the combined datasets consistently place Rbg1 at the interface of mRNA translation, nascent-chain maturation, and proteostasis.

It’s important to note that since nuclear proteins are synthesized on cytoplasmic ribosomes prior to nuclear import, changes in nuclear or mitochondrial pathways upon loss of Drg1 reflect changes in Rbg1/Drg1-mediated translation and do not necessarily indicate a nuclear site of Rbg1/Drg1 function.

### Lack of Rbg/Drg GTPases leads to delayed lag-to-exponential growth phase transition

Our growth analyses show that deletion of Rbg/Drg GTPase prolongs the lag phase and delays the transition from lag to exponential growth. This observation suggests that the function of Obg/Rbg/Drg GTPases likely extends to ribosomes containing unoccupied or deacylated tRNAs at the ribosomal A site under normal physiological conditions. During the acceleration phase of growth, when nutrient availability is high and ribosome biogenesis intensifies, large numbers of ribosomes and tRNAs are synthesized while aminoacyl-tRNA synthetases remain limiting. Under these conditions, uncharged tRNAs are likely to transiently occupy the ribosomal A site, increasing the likelihood of inadvertent activation of stress-response pathways, such as the stringent response in bacteria or the integrated stress response in eukaryotes. Our data suggest that Obg/Rbg/Drg GTPases protect this critical growth transition by maintaining translational homeostasis, ensuring that physiological pauses do not escalate into stress signals. The universal presence of a lag phase in growing cells underscores the fundamental need for controlled translational adaptation between quiescent and active states, suggesting the physiological importance of GTPase-mediated stabilization of translation during rapid biosynthetic acceleration.

### A model for Obg/Rbg/Drg-mediated protection of productive translation

Our results support a model in which Obg/Rbg/Drg GTPases recognize ribosomes experiencing increased translational or nascent-chain-associated demands and stabilize a state that remains competent for peptide synthesis. In this model, a slowed ribosome initially represents a potentially productive intermediate rather than an irreversibly stalled complex. Recruitment of Obg/Rbg/Drg, protects the ribosome, promotes peptidyl-transfer competence and allows additional time for tRNA accommodation, nascent-chain folding, targeting, or resolution of the underlying impediment. If the pause cannot be resolved, the ribosome may subsequently enter collision-dependent stress signaling or quality-control pathways. In this scenario, Rbg/Drg GTPase is recruited to slowed or partially stalled ribosomes in the translation cycle, including initiation, elongation, and termination/recycling. This GTPase is only able to promote a catalytic-competent conformation of the ribosome when the pause is physiological and temporary. In contrast, more severe or pathological stalls would persist beyond Rbg/Drg intervention, leading to sustained ribosome pausing or collisions that subsequently channel the complexes into alternative quality control pathways. In this way, Rbg/Drg may function as a key discriminator between temporary, physiologically meaningful pauses and harmful, persistent stalls.

This model distinguishes Obg/Rbg/Drg from canonical elongation factors that act during nearly every translation cycle. Obg/Rbg/Drg proteins may instead function selectively, increasing the probability of successful elongation when the ribosome occupies vulnerable intermediate states. Such a role explains why loss of Rbg1/Drg1 causes modest average effects in reporter assays while potentially exerting substantial consequences on specific endogenous proteins and cellular pathways.

Using this model, we hypothesize that, in bacteria, the monomeric Obg-like NTPases such as CgtAE/ObgE (48) or YchF act directly on translating ribosomes to prevent premature engagement of RelA and activation of the stringent response. In eukaryotes, the heterodimeric Rbg1/Tma46 in yeast (and Drg1/Dfrp1 complex in human) fulfills an analogous role by protecting paused ribosomes from stress-activated factors such as Gcn2. Despite their structural divergence, these GTPases share a conserved ribosome-binding G domain architecture that allows them to interact with the GTPase-associated center and modulate the ribosomal conformational dynamics. This evolutionarily conserved feature suggests that the ability to maintain ribosomal competence during transient translation pauses arose early and was preserved as a fundamental regulatory feature of the translational apparatus.

The structural evidence, in particular, supports our model in which ribosomal recruitment of Drg GTPases prevents premature activation of stress-response pathways. Based on the cryoEM structures of an Rbg1/Tma46-bound ribosome, when the ribosome encounters a translational pause, the E-site tRNA dissociates, and the head of the 40S subunit pivots, creating an enlarged gap between the head and shoulder domains. The conserved zinc-finger domain of Tma46 (Dfrp1 in human) inserts into this gap, and the GTPase complex Rbg1/Tma46 stabilizes the paused ribosome in a translationally competent conformation (3). Moreover, we also showed that the presence of yeast Rbg1 increases the stability of reporter mRNAs harboring a strong stalling sequence (3), suggesting that continued translation by Rbg GTPase diverts stalled translation complexes away from quality control pathways that degrade the affected mRNA.

Presence of Gcn1 in our RNase-treated IP ribosomal samples, combined with the structural evidence of Gcn1-bound collided disome (49), suggests that Rbg/Drg GTPase also binds to the collided ribosomes, in addition to a paused or stalled monosome. Structural comparisons show that Rbg/Drg binding sterically precludes Gcn2 association with the same ribosome. Because Gcn2–ribosome interaction is required for activation of the integrated stress response (ISR) (36–38), the presence of Rbg/Drg on paused or collided ribosomes likely serves to shield them from inadvertent stress signaling. Consistent with this interpretation, Gcn2 was not detected in any Rbg1 immunoprecipitated ribosomal samples, supporting the idea that Rbg/Drg binding and Gcn2 recruitment are mutually exclusive events.

Earlier genetic evidence also provides support for this model. It was shown that one of the yeast Rbg GTPases, Rbg1 or Rbg2, is required for optimal growth in yeast lacking the Slh1 gene (34), suggesting that Rbg proteins and Slh1 are most likely to function in different cellular pathways that have related functions. The protein product of this gene, Slh1 protein, is an ATP-dependent helicase that dissociates the two subunits of a stalled leading ribosome in a collided disome to commit it to ribosome-associated quality control (ASCC3 is the human counterpart of yeast Slh1) (50–54). Therefore, Rbg/Drg proteins likely play a role that is mechanistically different but genetically related to protein synthesis and quality control. Consistent with this prediction, we have not observed peptides belonging to Slh1 in Rbg1-IP-LC-MS/MS.

Obg/Drg is a relatively slow GTPase that would be excluded from the ribosome during the normal course of protein production; otherwise, the canonical translational GTPases, such as EF-G and EF-Tu (eEF-1 and eEF-2 in eukaryotic cells, respectively), would not be able to bind and function normally. In other words, Drg promotes translation when ribosomes pause, but the protein must leave the ribosome once its function is completed; otherwise Drg would instead become a translation inhibitor. Given the lack of rescue by the GTPase-deficient proteins in both the cell growth and *in vivo* translation assays (**Figure 2 to 5**), GTP-hydrolysis in Rbg/Drg likely at least facilitates its own disassociation from the ribosome, similarly to other translational GTPases such as EF-Tu and EF-G.

It is important to note that the WT ObgE stimulates puromycin incorporation into the K20-stalled ribosomal complex (**Figure 5**), whereas the D213G mutant inhibits the same reaction. The ObgE (D213G) mutant protein is capable of GTP-binding (the *in vitro* reaction buffer contains a high amount of GTP, so does the inside of the cell) but is incapable of hydrolyzing GTP. These results support that the process of GTP-hydrolysis in Obg facilitates conformational rearrangement in the PTC of the bound ribosome and promotes peptide bond formation. In the physiological text, because the factors that give rise to ribosomal pauses during translation are heterogeneous, as observed by our earlier genomic analyses (3), the GTP hydrolysis activity of the Obg/Rbg GTPase may facilitate ribosome progression across a range of conformational and energetic barriers associated with diverse pause signals. Through this mechanism, Obg/Rbg could help ribosomes and tRNAs navigate those relatively transient translational obstacles and thereby maintain efficient protein synthesis.

As mentioned earlier, several chaperones and co-chaperones that facilitate nascent chain folding, including Zuo1, Ssz1, Egd2, Ssb1 and Ssb2, Sti1 are enriched in Rbg1-immunoprecipitated ribosomal fractions, suggesting a close functional connection between nascent chain folding events and the rate of ribosome movement. Zuo1 and Ssz1 facilitate co-translational folding cooperatively with Ssb chaperones (43, 44). Egd1 and Egd2 are the components of the nascent polypeptide-associated complex (NAC), which regulates the specificity of co-translational ER targeting of ribosomes and likely help shield nascent peptides from co-translational ubiquitination (55, 56). Given these observations, *it is conceivable that the rate of protein synthesis on the ribosome is tightly monitored in the cell, both to ensure proper folding and to coordinate downstream events of targeting and quality control.* In this regard, our findings are reminiscent of an early observation in *E. coli* where exposure to chloramphenicol (which slows down ribosome movement) or streptomycin (which increases ribosome movement) induced the expression of cold-shock or heat-shock proteins, respectively (57).

The proposed model also accommodates evolutionary diversification. Bacterial Obg and eukaryotic Rbg/Drg need not possess identical interaction networks or physiological targets. Eukaryotic proteins have acquired partners and operate in a cellular environment containing more elaborate chaperone, targeting, signaling, and ribosome-quality-control systems. What appears to be conserved is the core capacity to support productive nascent-chain synthesis at challenged ribosomes. Lineage-specific proteins and domains may adapt this ancestral activity to different cellular contexts.

## Conclusions and future directions

Together, the phylogenetic, genetic, biochemical, transcriptomic, and proteomic results identify Obg/Rbg/Drg as an ancient GTPase pathway that supports productive translation. The data obtained thus far suggests that Obg/Rbg/Drg proteins act as protectors and facilitators of productive pausing. A pause does not necessarily represent irreversible damage; it can provide time for tRNA accommodation, nascent-chain folding, protein targeting, or cellular regulation. Obg/Rbg/Drg GTPases may help protect the ribosome by excluding the initiation of quality control on the ribosome and preserving the catalytic competence of these paused ribosomes, thereby preventing premature conversion of a recoverable pause into a pathological stall.

What Rbg/Drg GTPase does in the cell is an unfolding puzzle. Its name, Drg in human, originated as a historical accident, and is likely to be quite a misnomer. Given its direct association with the central translation machinery, ribosomes, and its function in enhancing protein biosynthesis upon ribosome pauses, calling it “developmentally regulated GTP-binding protein” thus underestimates the influence and depth of this protein in regulating gene expression. Future studies should determine the kinetic step affected by Obg/Rbg/Drg, define the ribosomal conformation recognized by these proteins, and establish how GTP binding and hydrolysis regulate association and release. Selective ribosome profiling and nascent-proteome analyses will be important for distinguishing Rbg1-associated messages from transcripts whose translation is functionally dependent on Rbg1. It will also be important to determine how Rbg1/Drg1 activity intersects with Gcn1/GCN2 signaling, collision recognition, chaperone recruitment, and ribosome-associated quality control. Additionally, possible nuclear functions of Rbg/Drg GTPases remain to be uncovered.

Overall, our findings suggest that cells possess a conserved mechanism that protects ribosomes before translational difficulty becomes irreversible. By maintaining the catalytic competence of challenged ribosomes and coordinating translation with nascent-chain proteostasis, Obg/Rbg/Drg GTPases may help determine whether a ribosomal pause remains productive or progresses toward pathological arrest. Investigations into the conserved regulatory roles these GTPases serve in cells across the kingdoms of life, with emphasis on how they impact physiological pathways, are critical to advancing our knowledge of how protein synthesis is integrated into the complex network of cellular gene expression.

## Acknowledgments

We thank the proteomics cores at UIUC and the University of Chicago for mass spectrometry, UIUC Carl R. Woese Institute for Genomic Biology for imaging, and Roy J. Carver Biotechnology Center for high-throughput RNA sequencing. We thank Dr. V. Ramakrishnan and Dr. Vish Chandrasekaran for sharing the mammalian K20 mRNA construct, Dr. William Metcalf for the archaeal strain, Dr. Albert Weixlbaumer and members of the Jin Lab for many helpful discussions. This research was funded by the startup offered to H. Jin from the University of Illinois at Urbana-Champaign and the National Science Foundation (Award Number: 2408763) to H. Jin.

## Materials and Methods

### Phylogenetic Analysis

Protein sequences of Obg-related GTPases (Obg, ObgC and ObgM), DRG/Rbg-related GTPases (Drg/Rbg, Drg1/Rbg1, and Drg2/Rbg2), and Nog1 were retrieved from the NCBI Protein database on August 31^st^, 2026 (1). Sequences were selected from representative organisms across Archaea, Bacteria and Eukarya (2–4). Due to the polyonymous nature of GTPase nomenclature across organisms and literature, synonymous protein names are listed below in Table M1. Protein sequences within each GTPase family were first aligned separately using Constraint-based Multiple Alignment Tool (COBALT) (5). An initial maximum-likelihood phylogenetic tree was generated for each GTPase family using FastTree (6) using 1,000 bootstrap replicates. Representative sequence among isoforms were selected based on the longest length (7). Sequence artifacts such as those with long branching, and poor quality were removed by visual inspection (6). The sequences retained across all the GTPases families were combined and aligned using COBALT (5). Regions that contained gaps in columns in the majority of sequences were excluded. The final maximum-likelihood phylogeny was constructed using IQ-TREE v2 (8) with 1000 replicates of both ultrafast bootstrap (UFBoot) and SH-aLRT (Shimodaira-Hasegawa approximate likelihood ratio test). The resulting phylogeny was visualized and annotated in FigTree v1.4.4 (9). The accession numbers of the final sequences used are listed in Table M2.

**Table M1.** Alternative names of GTPases. All eukaryotic homologues of the bacterial ObgM are group under ObgM as part of this analysis.

| GTPase Name | Synonyms |
| --- | --- |
| Obg | ObgE/CgtAE, YhbZ, CgtA, BsObg |
| ObgC | ObgC/ ObgA-1/ ObgA-2/ AtObgC/ ObgC1/ ObgC2 |
| ObgM | ObgH1/ MTG2/ GTPBP5 |
| Drg | Rbg |
| Drg1 | Rbg1/ NEDD3 |
| Drg2 | Drg2/ Rbg2 |
| Nog1 | GTPBP4 |

**Table M2.**
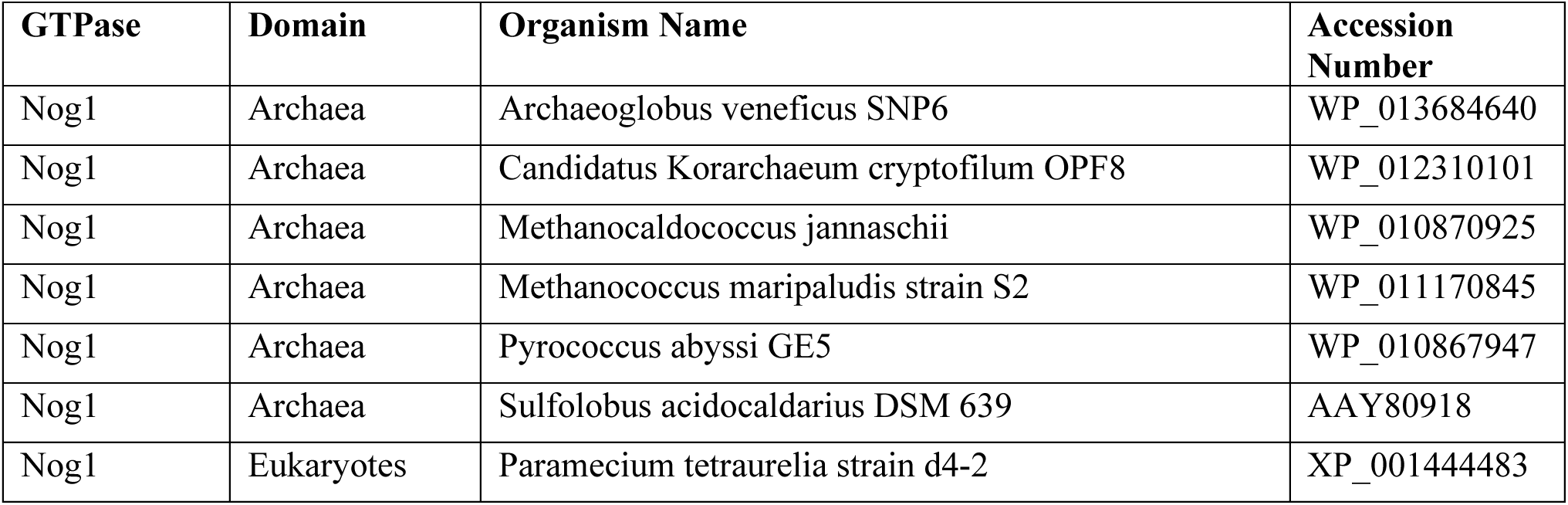
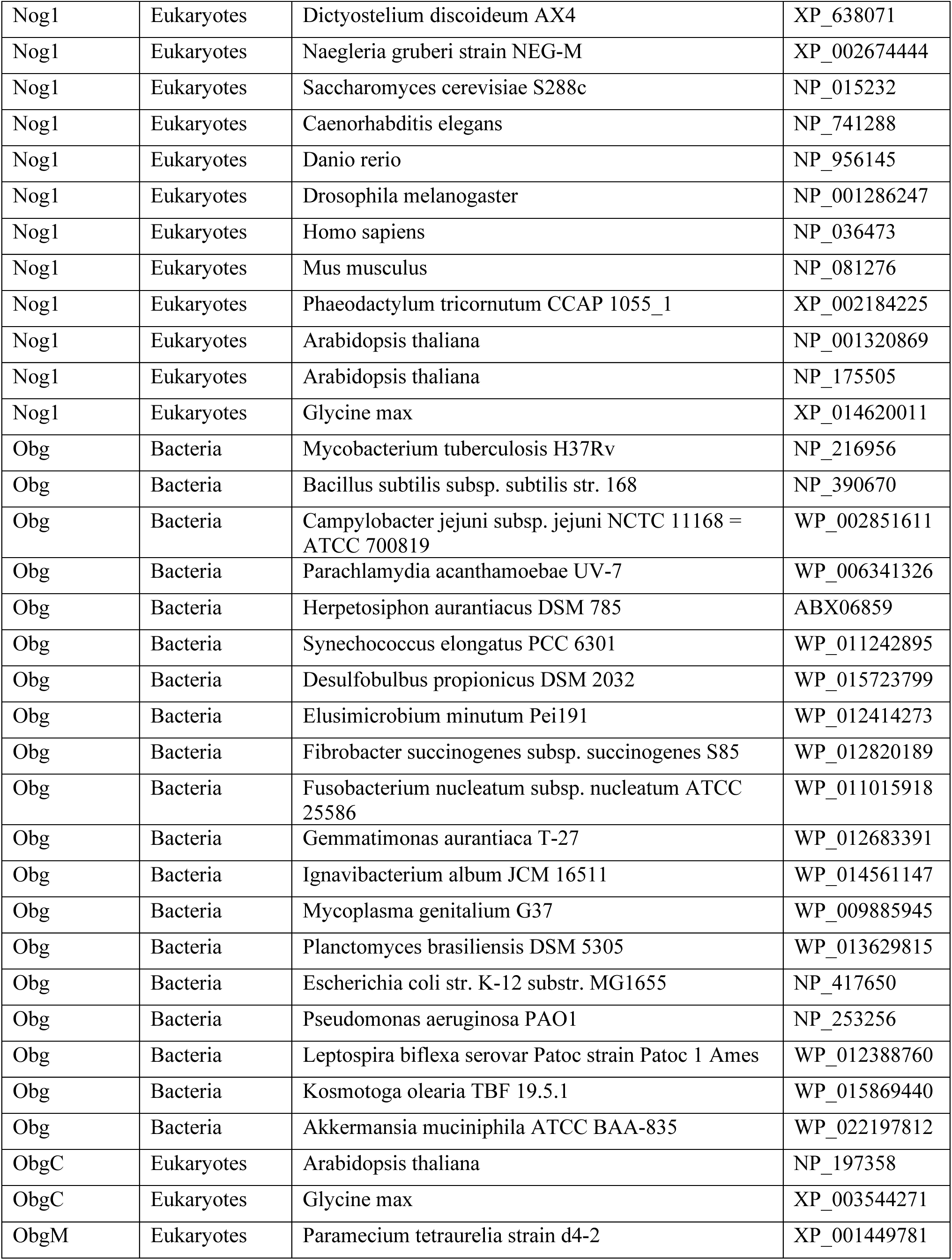
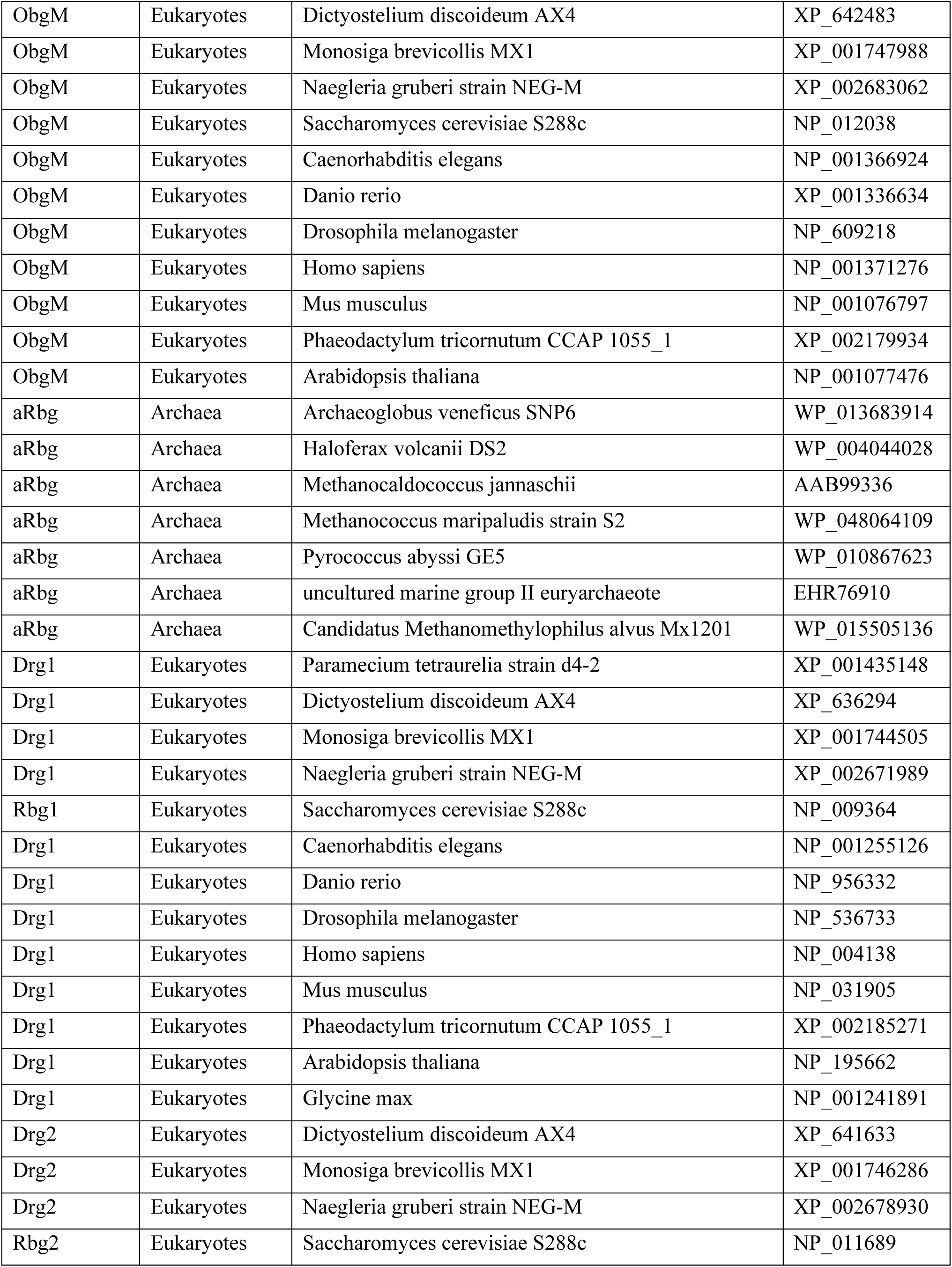
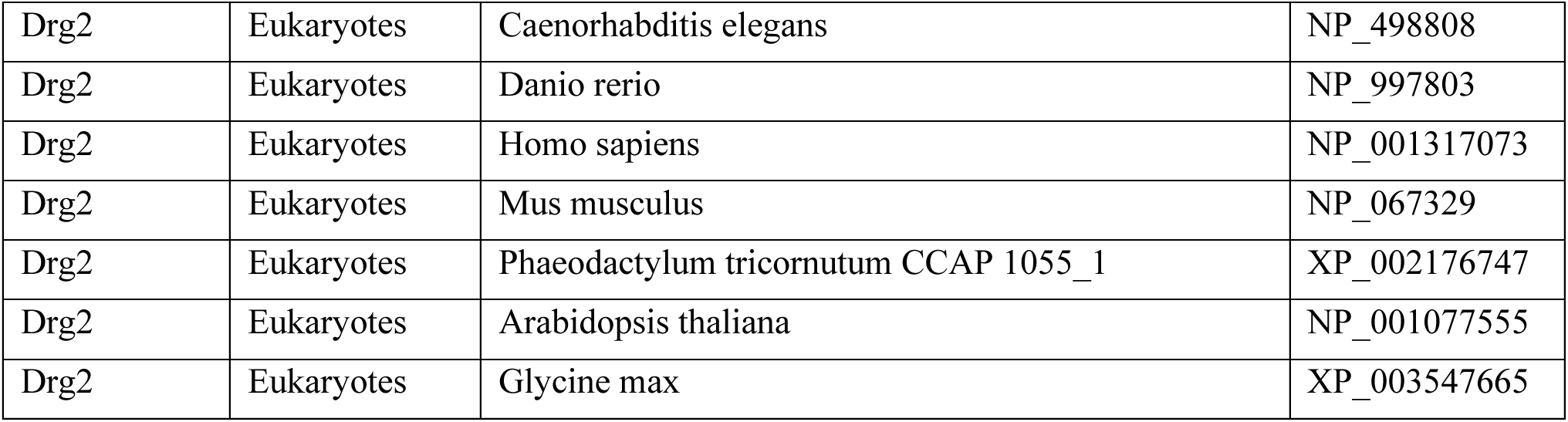
List of Accession Numbers Used in Phylogenetic Analysis. The table lists the GTPase, domain and organism name of each accession number used in the final phylogeny.

### Cell lines, strains, and growth conditions

The BY4742 strain of *Saccharomyces cerevisiae* (*MATα his3Δ1 leu2Δ0 lys2Δ0 ura3Δ0)* was used in this study. Yeast chromosomal knockouts were previously generated using homologous recombination techniques (10–12). Δ*rbg1* BY4742 *S. cerevisiae* used in this study has the following genotype: *MATα his3Δ1 leu2Δ0 lys2Δ0 ura3Δ0 rbg1::KanMX*, and was grown in YPD+G418 medium unless indicated otherwise. In addition, a BY4742 strain in which the endogenous *RBG1* locus was modified to encode a C-terminal FLAG tag with a TEV linker was used in this study. This chromosomally tagged *RBG1*-FLAG strain was selected on SC-Ura medium. When indicated, *S. cerevisiae* was transformed with recombinant vectors via the established LiAc/ssDNA/PEG method, with minor modifications (13).

Adherent HEK293T cells were used in this study. They were cultured in DMEM medium supplemented with 10% FBS, 1 mM Pyruvate, and 100 U/mL Penicillin + 100 μg/mL Streptomycin and grown in a 37°C incubator containing 5% CO_2_. See the separate methodology section for information regarding HEK 293T *Drg1* genomic knockouts.

### Drg1 Knockout in HEK 293T cells using CRISPR-Cas9

#### *In vitro* transcription of gRNAs

Single-guide RNAs (sgRNAs) targeting Exons 2 and 3 of the *Drg1* gene were synthesized by assembly PCR and *in vitro* transcription, as previously described (14). Briefly, a T7 promoter-containing template was generated using two partially overlapping primers (T7F_DRG1 and T7RevLong, 10 nM each) and flanking amplification primers (T7FwdAmp and T7RevAmp, 200 nM each), followed by 33 PCR cycles with Phusion High-Fidelity DNA Polymerase (NEB, M0531S) (Table M1). The PCR product (8 µL) was transcribed using the HiScribe T7 High Yield RNA Synthesis Kit (NEB, E2040S), followed by DNase I digestion (1 µL, 15 min at 37 °C). Transcribed sgRNAs were then treated with calf intestinal phosphatase (CIP; 2 U/µL, NEB, M0290S) in NEBuffer 2.1 (NEB) at 37°C for 3 h in a final volume of 100 µL. gRNAs were purified using the RNA Clean & Concentrator-5 Kit (Zymo Research, R1013) per the manufacturer’s protocol.

**Table M1:**
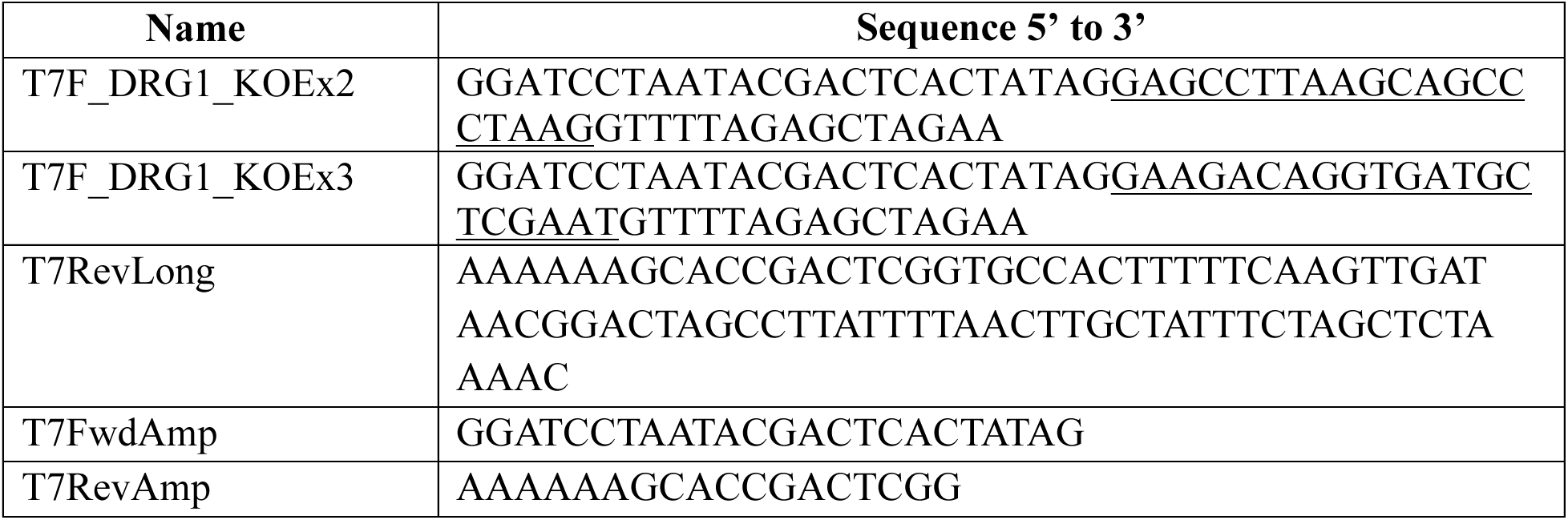
Primers used for PCR synthesis of sgRNA-encoding DNA templates for T7 polymerase-mediated *in vitro* transcription.

#### Transfection

Cells were seeded in 6-well plates to reach approximately 60-70% confluency at the time of transfection. RNP complexes were formed by incubating 0.5 µL of each gRNA (DRG1_KOEx2, DRG1_KOEx3) with 1.4 µL Cas9-GFP-Pro (Sigma, CAS9GFPPRO-50UG) in 250 µL OptiMEM® for 10 min, followed by the addition of 5 µL TransIT-X2® reagent (MirusBio) with an incubation for 15 min. A second transfection using the same procedure was performed the following day. After 24 h, 100 µL of cell suspension (5 cells/mL) was plated into individual wells of 96-well plates. After two weeks, single colonies were expanded into 6-well plates for further analysis.

#### Knockout analysis

Monoclonal populations were lysed in 30 µL lysis buffer (150 mM NaCl, 1% NP-40, 50 mM Tris-HCl, pH 8.0), resolved on a 12% SDS-PAGE gel, transferred to a nitrocellulose membrane, and probed with anti-DRG1 (Abcam, ab133648) followed by an HRP-conjugated anti-rabbit secondary antibody (Abcam, ab97051). Detection was performed using a BioRad ChemiDoc MP system. To confirm the knockout by sequencing, genomic DNA was extracted by heating cell pellets in 100 µL EB buffer (10 mM Tris-Cl, pH 8.5) at 95 °C for 15 min. Target regions flanking Exons 2 and 3 of *Drg1* were amplified using Phusion DNA polymerase (NEB, M0531S) with the following primers: (EX2_DRG1_F – 5’-GCCCTGAGAGCGCTTTACAATTAG-3’; EX2_DRG1_R – 5’-CACTGCACCCAGCGCATGTAAATC-3’) and (DRG1_EX3_Kn_F – 5’GTGCCTAGAGGGATAGGCAGTA-3’; DRG1_EX3_Kn_R – 5’-TTTGTATCTGATGACACCAGGC-3’). Amplicons were purified and analyzed by Sanger sequencing. To test the knockout clone for all alleles, Cas9 cleavage was assayed on an Exon 2 PCR product. Reactions (30 µL) contained 300 nM sgRNA, 2 µL Cas9 nuclease (1 µM), 3 µL NEBuffer 3 were pre-incubated for 10 min at room temperature. Subsequently, 3 µL target DNA (30 nM) was added and incubated at 37 °C for 1 h. Proteinase K (1 µL, 20 mg/mL) was then added, and samples were incubated for 10 min at room temperature. Products were analyzed on 1% agarose gel.

### Anisomycin cell viability assay

WT (wild type) and *Δrbg1* BY4742 cells were transformed with plasmids encoding Drg proteins from different species and *E. coli* EF-Tu that were expressed under a PGK promoter on a pJK372 vector. Successfully transformed pJK372 plasmids in BY4742 were first selected in uracil-lacking medium (-ura) SC-Ura medium (MP Biomedicals), as the URA3 gene is encoded by the pJK372 plasmid. Transformed cells were grown to the mid-log phase at 30 °C with shaking in SC-Ura medium (MP Biomedicals). Cell growth medium was switched to YPD after the initial selection to assay cell growth without the confounding effects of auxotrophic selection pressure. WT cells were grown in YPD medium. Different amounts of anisomycin were added to YPD to assay the cell growth behavior in the presence of anisomycin (MedChemExpress).

A yeast spot assay using serial dilutions was performed with modifications (15). Fresh selective medium was inoculated with cells to an OD_600_ value of 0.2, then the cells were grown until mid-log phase (approximately OD_600_ 0.6). Cells were harvested by centrifugation and resuspended at an OD_600_ of 1.0 in 1X PBS. Serial dilutions were made as indicated, in which 1 μL of each dilution was spotted on YPD, YPD+10 μg/mL anisomycin, and YPD+12.5 μg/mL anisomycin plates. The plates were grown at 30°C for the indicated times. Two replicates of the YPD and YPD+10 μg/mL anisomycin spot assay were completed, with one appearing in Fig. 1A and the other in Supplementary Fig. 2.

For the growth curves shown in Fig. 1, the indicated transformed and untransformed cells were first selected using auxotrophic selection pressure as described above and then collected by centrifuging at 6000 RCF for 5 min at room temperature. Their growth medium was changed to YPD medium, either containing or lacking the indicated concentration of anisomycin (MedChemExpress). Cells were resuspended to a cellular density of an optical density (OD_600_) of 0.05. 200 μL of each diluted culture was added to Greiner clear flat-bottom 96-well plates. Plates were incubated at 30°C in a POLARstar Omega plate reader, which took OD_600_ measurements at 30 min intervals over the course of the assays. Nine biological replicates were shown for the assay containing 7.5 μg/mL anisomycin, eight replicates were shown for the assay without anisomycin, and three replicates were shown for the assays containing 0.5 and 1 μg/mL anisomycin. Data points and error bars indicate the mean OD_600_ reading for all replicates and the standard error of the means, respectively. Log-phase doubling times were calculated by fitting a logarithmic regression to the exponential phase of the growth curves. The same procedure was used to study the growth behavior of the cells under stress, including growing at an elevated temperature of 37°C and in the presence of different levels of anisomycin at the elevated growth temperature.

### Alamar Blue assay

5,000 HEK293T cells were seeded per well in one row of a 96-well plate, and 5,000 DRG1-deficient (Δdrg1) HEK293T cells were seeded per well in five additional rows. A row containing complete DMEM supplemented with 10% fetal bovine serum and penicillin–streptomycin, but no cells, served as the blank. Twenty-four hours after plating, four rows of Δdrg1 cells were separately transfected with 200ng/ul plasmids encoding full-length human Drg1, the GTPase-deficient hDrg1(S78N) mutant, full-length bacterial ObgE, and the GTPase-deficient ObgE(S173N) mutant. The transfection was done using the TransIT reagent and Opti-MEM. The remaining Δdrg1 row was left untransfected. Twenty-four hours after transfection, Alamar Blue reagent was added, and cells were incubated for 3 hours. Then, the fluorescence was measured with an Agilent BioTek Cytation 5 Cell Imaging Multimode Reader at 570 nm excitation and 595 nm emission. The assay measures cellular reducing activity indicating cellular viability through conversion of resazurin to fluorescent resorufin. Blank-well fluorescence was subtracted from each experimental measurement, and the corrected values were plotted in GraphPad Prism.

### Protein expression and purification

BL21(DE3) cells were transformed by a standard heat shock method with a pET28-a vector expressing WT *E. coli* ObgE with an N-terminal 6X-His tag. ObgE mutants were generated by site-directed mutagenesis, verified by sequencing via Plasmidsaurus. Proteins were expressed in BL21(DE3) cells upon addition of 300 μM IPTG at 22°C. Cells were harvested, washed, and resuspended in 1X lysis buffer (20 mM Tris-HCl, pH 7.5; 500 mM NaCl; 1.5 mM MgCl_2_; 50 mM Imidazole; 1X Complete EDTA-free Protease Inhibitor (Roche); 40 U DNaseI (NEB)). After cell lysis, the lysates were clarified and incubated with Ni-NTA Agarose (Qiagen) at 4°C for 1.5 h. Afterwards, they were applied to the Ni-NTA column and washed with 1X lysis buffer lacking protease inhibitor and DNaseI. Protein was eluted using 1X elution buffer (20 mM Tris-HCl, pH 7.5; 500 mM NaCl; 1.5 mM MgCl_2_; 500 mM Imidazole), then concentrated with 30 kDa MWCO Amicon centrifugal filters (Millipore Sigma). Concentrated fractions were further purified via a Superdex 200 10/300 size-exclusion column in an AKTA FPLC (GE) in ObgE Storage Buffer (20 mM Tris-HCl, pH 7.5; 150 mM KCl; 5 mM MgCl_2_; 2 mM 2-mercaptoethanol), concentrated, and flash frozen in liquid nitrogen and stored at –80°C.

### *In vivo* flow cytometry translation assay

S. cerevisiae BY4742 and its Δrbg1 derivative were transformed with the LEU2-marked R12 dual-fluorescence reporter pJK372 plasmid alone (WT and Δrbg1 conditions), or co-transformed with the reporter plasmid and a second, URA3-marked pJK372 plasmid sharing the identical backbone and expressing Rbg1, Rbg1(S79N), ObgE, or ObgE(S173N), using a standard lithium acetate/PEG transformation protocol. For each condition, three independent transformants served as three independent biological replicates. Single colonies were picked from SC−Leu (reporter-alone strains) or SC−Leu−Ura (GTPase-expressing strains) selective plates and grown overnight at [30°C, 200 rpm] in the corresponding liquid selective medium, then diluted into fresh selective medium and grown to mid-log phase (OD₆₀₀ ≈ [0.5–0.6]) prior to analysis. Cells were washed in PBS and analyzed by flow cytometry equipped with FITC and TE, collecting ∼30000 singlet events per sample; a fluorescence intensity gate excluding events below 10² RFU in either channel was applied to exclude non-fluorescent/auto fluorescent cells.

Actively dividing WT and *Δdrg1* HEK293T cells were trypsinized and passaged in cell culture-treated 6-well dishes (Thermo) at a cellular density of 1×10^5^ cells/mL. After 24 h, cells were transfected with pEGFP-N1 plasmids encoding the dual-fluorescent reporter mRNA and, when indicated, modified pEGFP-N1 plasmids encoding Drg proteins on strong CMV promoters (or equimolar concentrations of corresponding empty vectors) using the Transit293 transfection system (Mirus Bio) according to the manufacturer’s recommendations. After 24 h transfection, cells were harvested and washed twice with ice-cold 1X PBS and stored in ice until measurement. A BD Symphony A1 flow cytometer was used to selectively excite and measure emission for both cellular RFP (mCherry) and GFP (EGFP). FCS Express 7 was used to gate and isolate events from individual cells. The violin plot was generated with the ggplot2 R package. *P*-values were calculated for the comparison of means of median values from three replicates using a unpaired two-tailed t-test. 23,000 individual cells were measured for each replicate. The replicates are reported in **Supplemental Figure 6.**

### Puromycin incorporation assay

*E. coli in vitro* translation was used to measure puromycin reactivity in stalled ribosomes. The experiment was conducted similarly to a previously published method using mammalian translation systems (16) with modifications. The vector encoding the 3’-truncated mRNA used to assay puromycin incorporation in ribosomes harboring polylysine tracks, was modified from a vector gifted by the Ramakrishnan laboratory of the MRC LMB in Cambridge. T7 promoter and Shine-Dalgarno sequences were added at the 5’ end and the 3’ end was truncated and modified to contain zero lysine (for the K0 construct) or twenty consecutive lysines by PCR (for the K20 (AAA) construct). In addition to the 3’ test sequences, the recombinant DNA encodes an N-terminal 3XFLAG tag, and a central villin headpiece domain (VHP). The mRNA construct was co-transcriptionally translated using the defined, reconstituted *E. coli* PURExpress ΔRF123 expression system (NEB) at 37°C for 1 h. Expression was halted by placing the reactions on ice. 2 μM ObgE or an equal volume of ObgE Storage Buffer and 2 μM puromycin (Sigma) were added to the reaction. Puromycin-treated translation reactions were returned to 37°C. Aliquots were taken at the indicated time points and quenched by diluting in SDS-PAGE loading buffer and incubating on ice. Puromycin-reacted and released peptides were separated from peptidyl tRNAs by SDS-PAGE and then visualized via anti-FLAG immunoblotting. For each data point, the percentage of reacted puromycin-conjugated peptides was quantified using ImageJ (NIH) by measuring the integrated density of the corresponding band, divided by the sum of both the puromycin-conjugated peptide band and unreacted peptidyl-tRNA band intensities. The data are represented as an average of at least 3 replicates with standard errors shown as error bars.

### SDS-PAGE and Immunoblotting

Protein samples were analyzed by standard SDS-PAGE and immunoblotting techniques. For routine SDS-PAGE, proteins were electrophoresed through 12% polyacrylamide (Biorad) gels running at a constant 160 V and stained with Coomassie blue. For routine immunoblotting for quantification, samples were quantified using Bradford Assay (Thermo), and an equal amount of total protein for each sample was loaded and SDS-PAGE was conducted at a constant 160V for approximately 1 h, followed by transfer to nitrocellulose membranes (Amersham) upon which M2 Anti-FLAG mouse antibodies (Sigma, 1:10,000 dilution) were used to visualize 3X-FLAG fusion peptides. For immunoblotting used for puromycin incorporation, NuPAGE 4-12% Bis-Tris gels (Thermo Fisher) were loaded with protein samples and SDS-PAGE was conducted at a constant 160V for approximately 1 h. Protein samples were transferred to nitrocellulose membranes (Amersham) upon which M2 Anti-FLAG mouse antibodies (Sigma, 1:10,000 dilution) were used to visualize 3X-FLAG fusion peptides.

### ObgE binding to the 80S ribosome analyzed by sucrose gradient

Δrbg1 BY4742 cells were transformed with vector expressing ObgE-FLAG under PGK promoter on a pJK372 vector and was selected under using auxotrophic selection pressure as described above in the methods section. A single colony was used for overnight culture under selection media (SC-URA) and was further grown in YPD to OD600 of 0.6. The cells were then treated with 100ug/mL of CHX for 5 mins and pelleted. The pelleted cells were resuspended and popcorned using lysis buffer (20mM Tris-HCl pH 7.5, 5mM MgCl_2_, 150mM KCl, 0.002mM ZnCl_2_, 0.5% Triton X-100, 100 U RNase Inhibitor, 1X Protease Inhibitor, 100ug/mL of CHX) in liquid nitrogen. To disassociate the two ribosomal subunits, 10mM of EDTA was added to the lysis buffer instead of CHX. The popcorned cells were lysed and the lysate was clarified by spinning at 4700 RPM for 10 mins at 4°C. The clarified lysate was used to measure A_260_ and subsequently an appropriate amount of lysate was used for sucrose gradient (10-50%) for profiling. The samples were spun for 32000 RPM in SW 32 TI swing bucket rotor for 3 hours 45 mins at 4°C using Beckman Coultier Ultracentrifuge. The samples were pulled and collected as depicted in the figure and the collected samples were then used for Western Blot using Anti-FLAG primary Antibody (Sigma) and Anti-Mouse secondary Antibody with HRP (Invitrogen). The blot was generated by incubating with ECL (Thermo) and imaged using iBright-1500.

### mRNA-seq

#### Sample Preparation

Saccharomyces cerevisiae BY4742 strain was used and FLAG-tag was inserted at the 3’ end of Rbg1 chromosomal locus by using established homologous recombination techniques. Chromosomally FLAG-tagged cells were grown at 30°C until they reached an OD600 of 0.6, and were treated with 100 µg/ml of cycloheximide for 5 minutes before harvesting. Cells were quickly harvested by centrifugation and frozen. The frozen cell pellet was ground under liquid nitrogen with 1 ml of 3x polysome lysis buffer supplemented with EDTA-free protease inhibitors (Roche), SUPERaseIN using an RNase-free mortar and pestle. Grounded samples were left on ice slurry to thaw and lysate was clarified at 18,000 × *g* for 10 minutes at 4°C, then loaded onto 10%–50% sucrose gradients in polysome buffer (20 mM Tris-HCl, pH 8.0, 140 mM KCl, 5 mM MgCl2, 20 U/ml SUPERase In (Fisher Scientific), 0.5 mM DTT, 100 μg/ml cycloheximide). Gradients were centrifuged at 32,000 rpm for 3 hours and 45 minutes in a Beckman SW 32.1 Ti rotor and polysome profiles were generated by fractionation with continuous measurement of absorbance at 260 nm. Fractions for 80S and polysome were pooled and the ribosomes were pelleted down by ultracentrifugation at 42,000 rpm for 15 hours in Type 45 Ti rotor. Pellets were resuspended in polysome gradient buffer and used for mRNA-seq.

For Rbg1-IP mRNA-seq library preparation, resuspended ribosomes were first incubated with 50 µL magnetic FLAG beads at 4°C for 2 hours and washed by polysome buffer for three times. RNAs were directly extracted from the beads with 500 µL Trizol. In parallel, regular total RNA was also isolated from the lysate, 80S and polysome pool. The extracted RNA was clarified and concentrated using Zymo kit (Zymo Research).

#### High-throughput Sequencing

Oligo-dT enrichment of RNA samples, quality control and construction of the RNAseq libraries and sequencing on the Illumina NovaSeq X Plus were performed at the Roy J. Carver Biotechnology Center at the University of Illinois at Urbana-Champaign. Purified DNase-treated total RNAs were run on a Fragment Analyzer (Agilent Technologies, Santa Clara, CA) to evaluate RNA integrity, followed by Oligo-dT enrichment. RNA-seq libraries were made using the Watchmaker mRNA Library Prep Kit (Watchmaker Genomics, Boulder, CO) according to the manufacturer’s protocol. Final libraries were quantified using a Qubit fluorometer (Thermo Fisher Scientific, Waltham, MA), and the average cDNA fragment size was determined using a Fragment Analyzer (Agilent Technologies, Santa Clara, CA). Libraries were diluted to 10 nM and further quantified by qPCR on a CFX Connect Real-Time PCR System (Bio-Rad Laboratories, Hercules, CA) to ensure accurate pooling of the barcoded libraries and maximize cluster density on the flow cell.

The pooled, barcoded RNA-seq libraries were loaded onto one NovaSeq X Plus 1.5B flow cell lane for cluster generation and sequencing (Illumina, San Diego, CA). Libraries were sequenced as paired-reads with 150 nt reads from each end. Raw base call files were converted to demultiplexed FASTQ files using bcl2fastq v2.20 Conversion Software (Illumina, San Diego, CA).

#### Data Analysis

Rbg1-associated mRNAs were defined by sequencing poly(A)-selected RNA recovered after immunoprecipitation of FLAG-tagged Rbg1 from pooled 80S and polysome fractions in glucose-grown cells. Matched lysate, polysome, and polysome-IP libraries from two biological replicates were processed through a common alignment and gene-counting workflow. Non-primary alignments and duplicate reads were removed, reads assigned to rRNA, snoRNA, tRNA, other noncoding RNA, and transposable-element features were excluded, and paired fragments overlapping annotated exons were quantified with featureCounts using Ensembl gene identifiers. Differential recovery was tested from raw integer counts with DESeq2. The paired design included replicate and fraction terms. Poly-IP-versus-polysome contrast measured Rbg1-specific recovery while controlling for the matched polysome pool. Algebraically, this contrast is equivalent to (Poly-IP/Lysate)/(Polysome/Lysate). DESeq2 median-of-ratios size factors normalized library depth, negative-binomial generalized linear models estimated log2 fold changes, Wald tests assessed significance, and Benjamini-Hochberg correction controlled the false-discovery rate. Poly-IP-enriched transcripts were defined by adjusted p < 0.05 and log2 fold change > 1, identifying 1,129 transcripts in the figure’s analysis universe. UniProt subcellular-location annotations were collapsed into Cytosol, Membrane, Endoplasmic Reticulum, Golgi, Mitochondrion, Nucleus, and Other classes. Sequence and protein features were compared between Poly-IP-enriched transcripts and the remaining tested pool. Transcript length and GC fraction were calculated from representative transcript sequences. A-rich codon fraction was the proportion of codons containing at least two adenines. Continuous features were compared using two-sided Mann-Whitney U tests with Benjamini-Hochberg adjustment and Cliff’s delta effect sizes. Gene Ontology over-representation was performed separately for Biological Process, Cellular Component, and Molecular Function with clusterProfiler, using all DESeq2-tested, ORF-mappable genes as the background and Benjamini-Hochberg-adjusted p values. Volcano plots displayed log2 fold changes and adjusted significance, while localization pie charts summarized the compartmental distribution of enriched protein products.

### Immunoprecipitation LC-MS/MS

BY4742 yeast encoding a C-terminal FLAG tag with a TEV linker on the chromosomal rbg1 gene were grown to mid-log phase (OD_600_ 0.6) in YPD medium, treated with 100 μg/mL cycloheximide for 5 min at 30℃ with shaking, harvested by centrifugation, resuspended in Lysis Buffer (20mM Tris-HCl, pH7.5; 150mM KCl; 5mM MgCl_2_; 2μM ZnCl_2_; 10U/mL SUPERase·In (Thermo; excluded in RNaseI treated samples); 100μg/mL cycloheximide (Sigma); 1X cOmplete EDTA-free protease inhibitor), then lysed by cryogenic grinding. Whole cell lysates were generated by centrifugation at 4000 rpm for 5 min, then cytosolic fractions were isolated after centrifugation of 14,000 rpm for 7 min. For RNase I treated samples, 10 U RNase I were added per 20 ug total RNA and incubated for 45 minutes at room temperature with gentle agitation. Cytosolic fractions were loaded on 2 mL 1M Sucrose cushions in the lysis buffer, then centrifuged at 39,000 rpm 4℃ for 3 h using Type 45 Ti Fixed-Angle Rotor. The ribosomal pellets were washed twice with wash buffer (20 mM Tris-HCl, pH 7.5; 150 mM KCl; 5 mM MgCl_2_; 2 μM ZnCl_2_), then resuspended in the wash buffer. For salt-wash control samples, resuspended pellets were subjected to 20 min end-over-end rotation incubation at 4°C, re-pelleted through 1 M sucrose cushion and resuspended in the wash Buffer. For immunoprecipitation (IP), samples were incubated with 40 μL of equilibrated αFLAG M2 Affinity Gel (Sigma) for 2 h at 4°C with gentle agitation. IP samples were washed four times with the wash Buffer, then digested on the beads as described in the following paragraph.

For Rbg1-IP-MS, which underwent a high salt wash (Supplemental Figure S8B), BY4742 cells individually overexpressing Rbg1-FLAG and Tma46-FLAG were cultured to OD_600_ of 0.6. The cells were then combined and popcorned using liquid nitrogen in lysis buffer (20mM Tris-HCl pH 7.5, 5mM MgCl2, 150mM KCl, 0.002mM ZnCl2, 0.5% Triton X-100, 1X Protease Inhibitor, 100ug/mL of CHX). The popcorned cells were lysed and the lysate was clarified by spinning at 4700 RPM for 10 mins at 4°C. The portion of clarified lysate was used for FLAG bead pull down and the remaining were treated with RNase-I and was subsequently used for sucrose gradient (10-50%) to collect the monosomes and spun for 3hours at 32000RPM at 4°C using SW 32 TI swing bucket rotor. The collected monosomes were further subjected to FLAG-bead pull down. After incubation with FLAG beads, the beads were washed with high salt, and elution was done using elution buffer (20mM Tris-HCl pH 7.5, 5mM MgCl2, 150mM KCl, 0.002mM ZnCl2) containing anti-FLAG peptide. The eluted samples were sent for mass spectrometry.

Proteomic analyses were performed at the Proteomics Core of the Roy J. Carver Biotechnology Center at the University of Illinois Urbana-Champaign. The samples were first reduced and alkylated. The samples were then digested with LysC followed by overnight trypsin digestions. The digested peptides were desalted with StageTips and then analyzed using a Q Exactive HF-X mass spectrometer (Thermo Fisher Scientific) coupled to an UltiMate 3000 UHPLC (Thermo Fisher Scientific) over a 60-minute reversed-phase gradient. The raw LC-MS data were searched against the Uniprot *Saccharomyces cerevisiae* reference proteome and user-provided sequence using Mascot v2.8.2. A reverse decoy database strategy was used to calculate the false discovery rate (FDR).

Protein quantification was performed using emPAI values exported from the Mascot search results. For each sample set, emPAI values were normalized to the total emPAI signal to account for loading and analytical variation. Differential protein abundance analysis was performed by calculating log_2_-transformed fold changes between experimental groups. Statistical significance was assessed using two-tailed t-tests, and proteins with *p* < 0.05 and absolute log_2_ fold-change > 1 were considered significantly enriched or depleted.

## Supplemental Figures and Figure Legends

**Supplemental Fig. 1.**
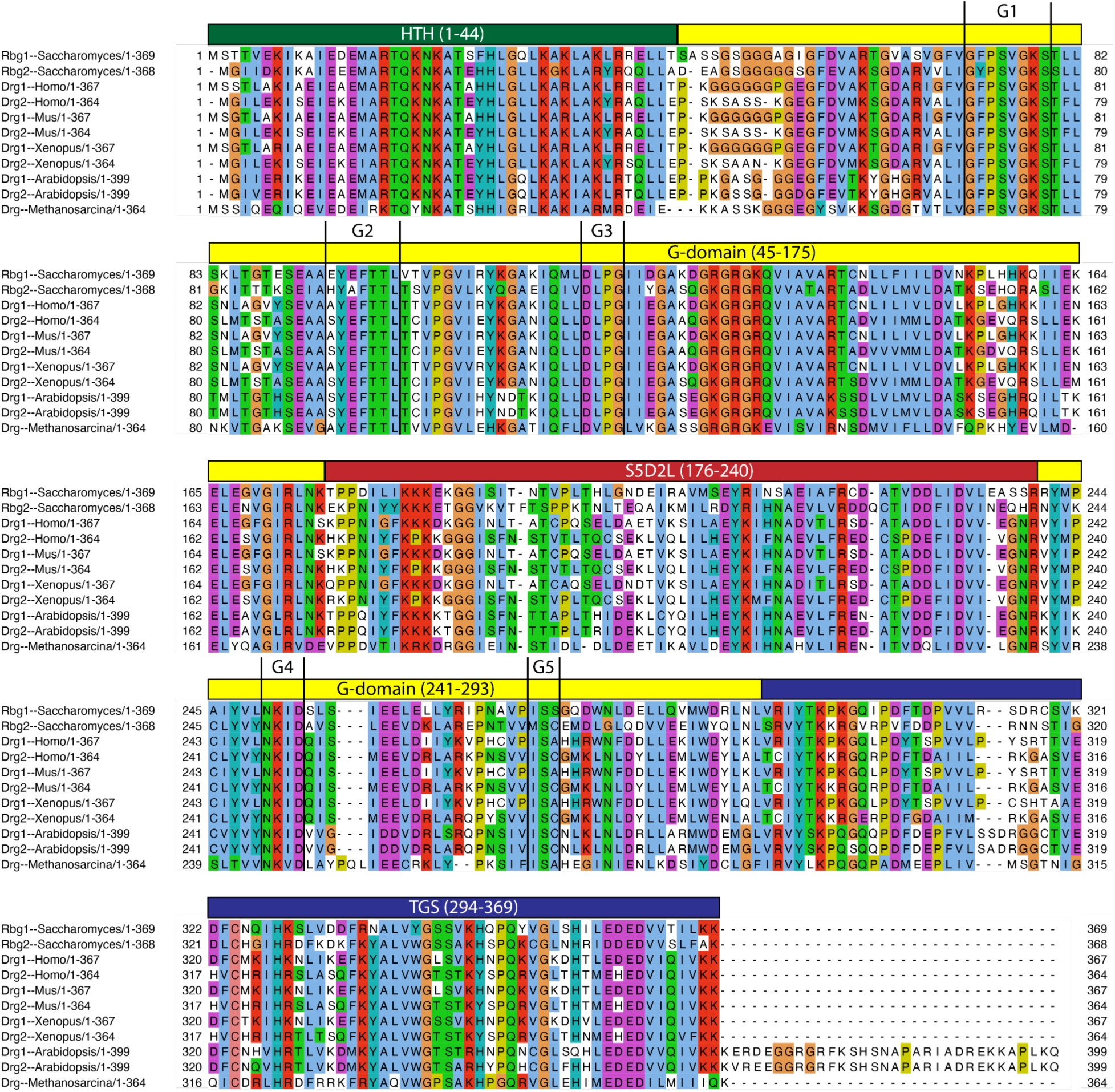
Amino acid sequences of Drg1 and Drg2 are highly homologous. Amino acid sequence alignment of yeast, human, mouse, *Xenopus*, *Arabidopsis*, and archaeal (*Methanosarcina*) Drg proteins with Rbg1 domain structure overlaid. The MUSCLE algorithm was used to generate the multiple sequence alignment. Jalview was used to visualize the multiple sequence alignment results. Residues are colored according to the Clustal color scheme in Jalview, in which a color is applied to each amino acid in the alignment only when the composition of residues at that position meets a defined percentage threshold specific to that residue category (Categories: Blue=hydrophobic, Red=positively charged, Magenta=negatively charged, Green=polar, Pink=cysteines, Orange=glycines, Yellow=prolines, Cyan=aromatic, White=unconserved).

**Supplemental Fig. 2.**
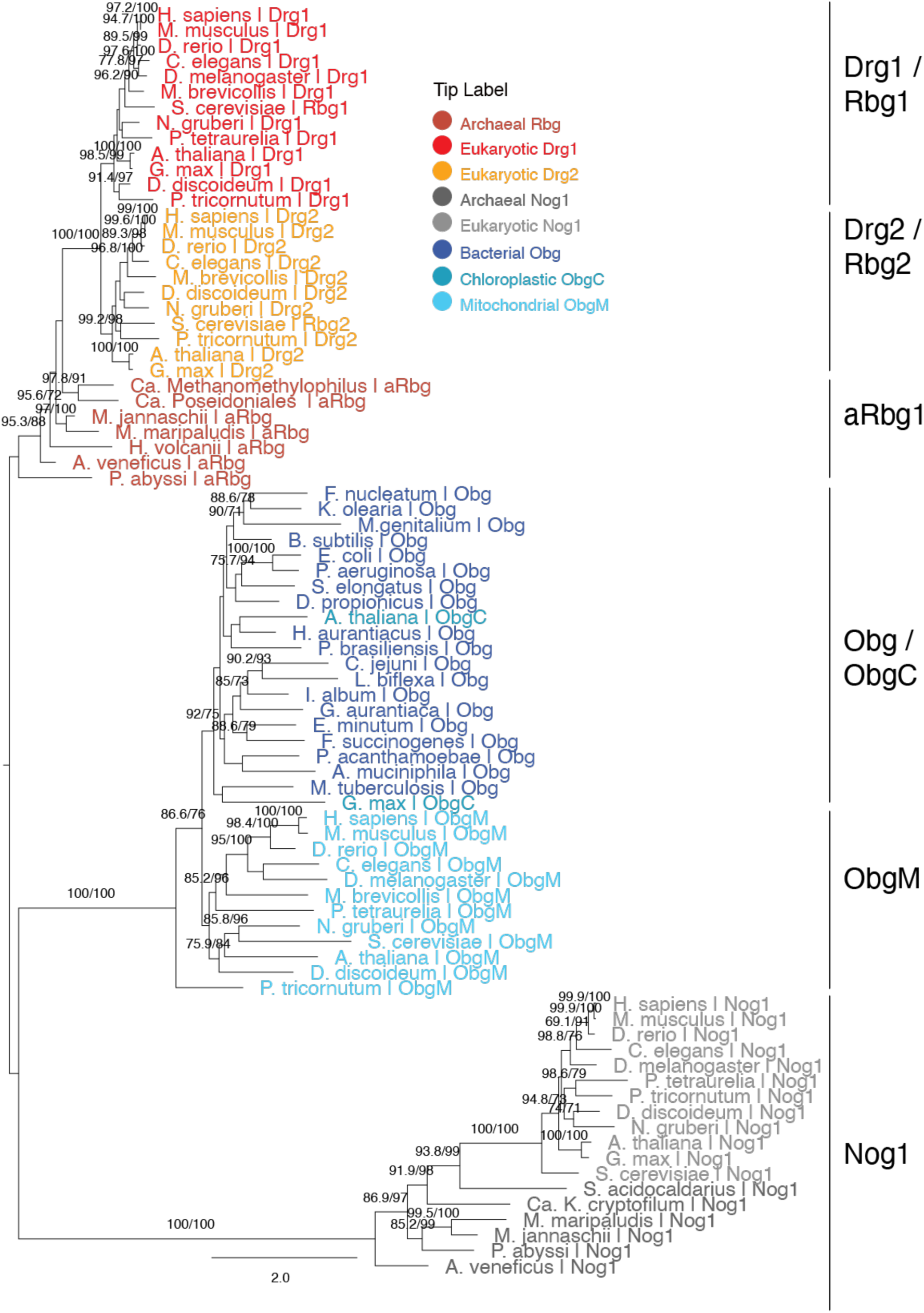
Phylogenetic relationships among Obg, Rbg/Drg and Nog1 family GTPases across the three domains of life. The phylogenetic tree was constructed from aligned protein sequences representing bacterial Obg proteins, archaeal and eukaryotic Rbg proteins, archaeal and eukaryotic Nog1 proteins, and eukaryotic organellar Obg homologs. Sequence labels specify the protein family. Detailed domain or major taxonomic group, organism, and accession number are included in the Methods section. Rbg-family sequences are shown in red, orange and brown, Obg-family sequences are shown in light blue, teal and dark blue, and Nog1-family sequences in light and dark grey. Numbers at internal nodes indicate branch-support values (SH-aLRT/UFBoot). Only SH-aLRT/UFBoot values greater than 70/70 are reflected. Branch lengths represent inferred evolutionary distances; the scale bar corresponds to 2.0 substitutions per site.

**Supplemental Fig. 3.**
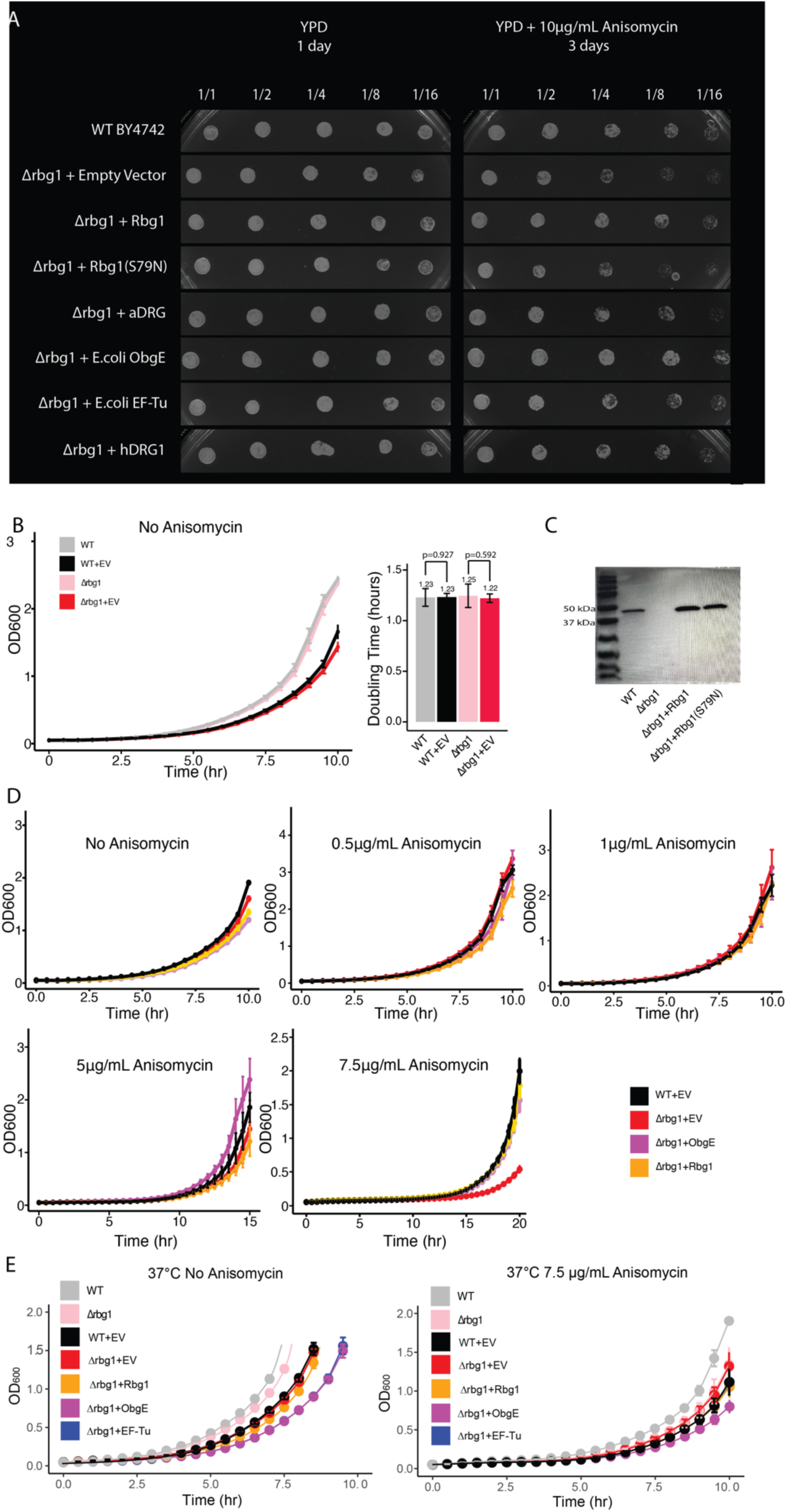
Drg orthologs rescue *Δrbg1* yeast growth in the presence of anisomycin. **A. Additional replicate of spot assay.** Serial dilutions of wild-type and *Δrbg1 S. cerevisiae* that were transformed with the indicated vectors were spotted on YPD plates with and without 10 µg/mL anisomycin and grown at 30 °C for the indicated times. **B. Plasmid transformation delays log-phase entry.** Wild-type and *Δrbg1 S. cerevisiae* transformed with empty pJK372 plasmids exhibit delayed log-phase entry compared to untransformed counterparts under standard growth conditions (YPD medium, 30°, no anisomycin). However, log-phase doubling times remain unchanged between transformed and untransformed strains. Data points represent mean A600 values from eight replicates. Error bars indicate standard error of the mean. **C. Relative expression of chromosomal and plasmid-expressed Rbg1 protein.** *a*FLAG immunoblot showing relative expression of chromosomally FLAG-tagged Rbg1 and plasmid-expressed (pJK372-PGK) Rbg1 or Rbg1(S79N) in *Δrbg1* yeast. **D. Dose-dependent rescue of *Δrbg1* yeast growth by plasmid-expressed Drg proteins** WT and *Δrbg1* yeast strains exhibit comparable growth in YPD at 0, 0.5, and 1µg/mL anisomycin. At 5 µg/mL anisomycin, growth defects emerge in *Δrbg1* strains and are partially rescued by ObgE expression. At 7.5 µg/mL, growth defects are more severe, but both ObgE and Rbg1 expression restore growth to near-WT levels. Data points represent mean A600 values at the indicated time points from eight replicates (0 µg/mL), three replicates (0.5, 1, and 5 µg/mL), and nine replicates (7.5 µg/mL). Error bars indicate the standard error of the mean. **E. A suitable amount of native Rbg1 is required for optimal growth under stress.** Growth behaviors of WT and *Δrbg1* cells were added to the data reported in Figure 1C for comparison of the growth of different cells under stress. All cells were monitored using optical density at OD_600_ over 10 hours at 37°C. Cells carried either an empty vector (EV) or a vector expressing the indicated protein (Rbg1, ObgE, or EF-Tu). Cell growth at 37°C without anisomycin (Left panel), and growth at 37°C with 7.5 μg/mL anisomycin. Data points represent the mean ± SEM of 8 biological replicates. The *Δrbg1* deletion strain exhibits a growth defect relative to WT under both conditions, which is rescued by re-expression of Rbg1 but not by the heterologous GTPases ObgE or EF-Tu. In the presence of anisomycin, WT cells display markedly enhanced growth relative to all *Δrbg1*-derived strains, indicating that the presence of a suitable amount of Rbg1 contributes to translational robustness under stress conditions.

**Supplemental Fig 4.**
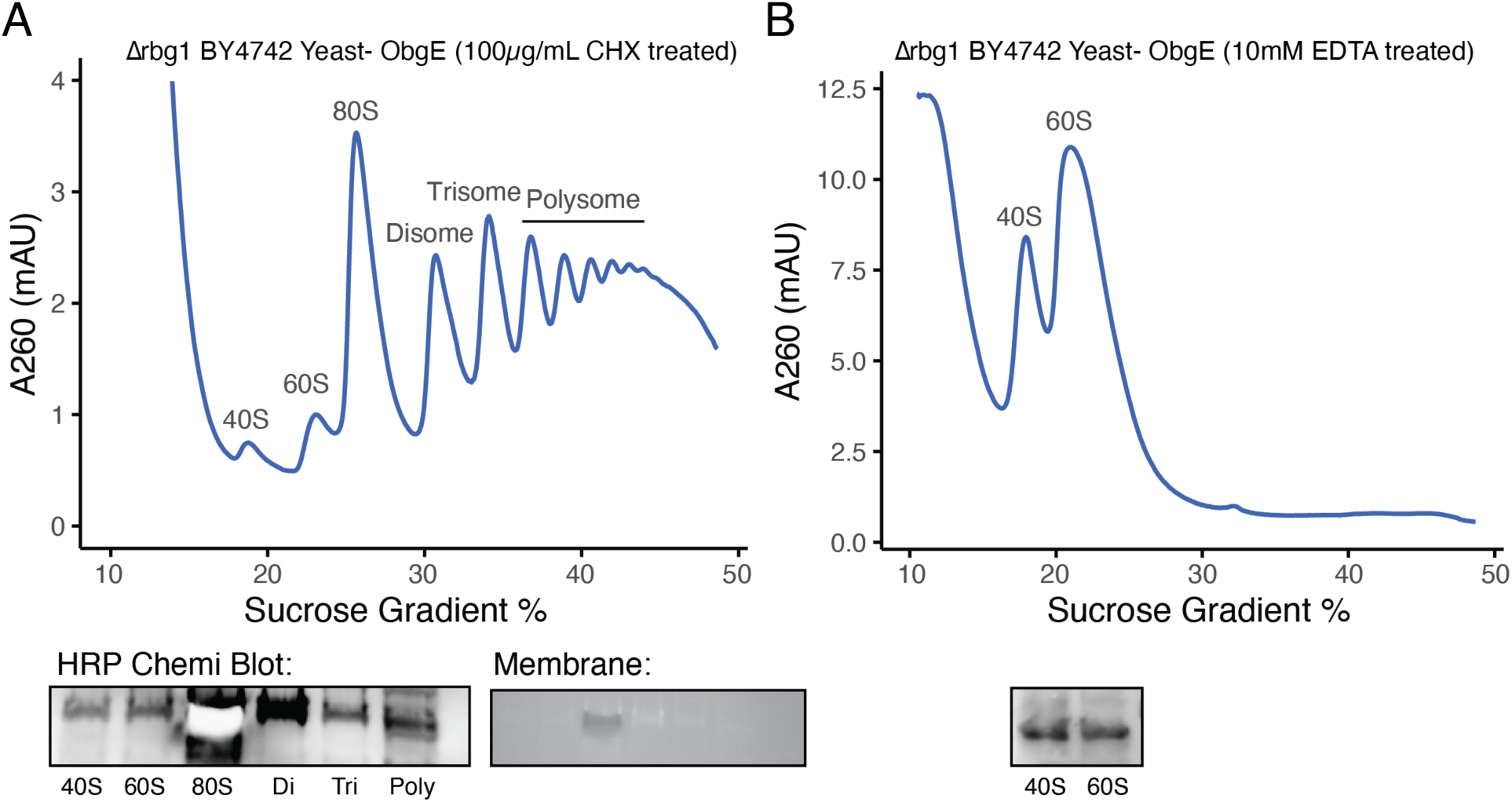
ObgE associates with 80S ribosomes in *Δrbg1* yeast cells. **A. Sucrose density gradient profile of *Δrbg1* BY4742 cells expressing E. Coli ObgE**. The polysome profile of lysates from *Δrbg1* BY4742 yeast cells expressing ObgE was obtained by sucrose density gradient sedimentation. The profile was monitored by absorbance at 260 nm (A260, mAU). Peaks of 40S and 60S ribosomal subunits, 80S monosomes, disomes, trisomes, and polysomes are indicated. Gradient fractions were collected and pooled. Western blot was performed. HRP chemiluminescence western blot and membrane are shown. ObgE is detected across translating ribosome fractions (disome through polysome) as well as in the 60S and 40S subunit fractions, suggesting that heterologously expressed ObgE associates with translating ribosomes in the *Δrbg1* yeast cells. Due to the presence of excess protein in 80S pool, the membrane got charred (as shown in only the membrane image) resulting a white blob like artifact in the HRP Chemi Blot image. **B. Sucrose density gradient sedimentation profile of lysates from *Δrbg1* BY4742 yeast cells expressing ObgE and treated with 10 mM of EDTA**. EDTA dissociates ribosomes into 40S and 60S subunits. Western blot of fractions corresponding to the 40S and 60S subunit peaks shows ObgE co-sediments with both dissociated subunits, suggesting ObgE bound to ribosomes followed by EDTA-induced dissociation.

**Supplemental Fig. 5.**
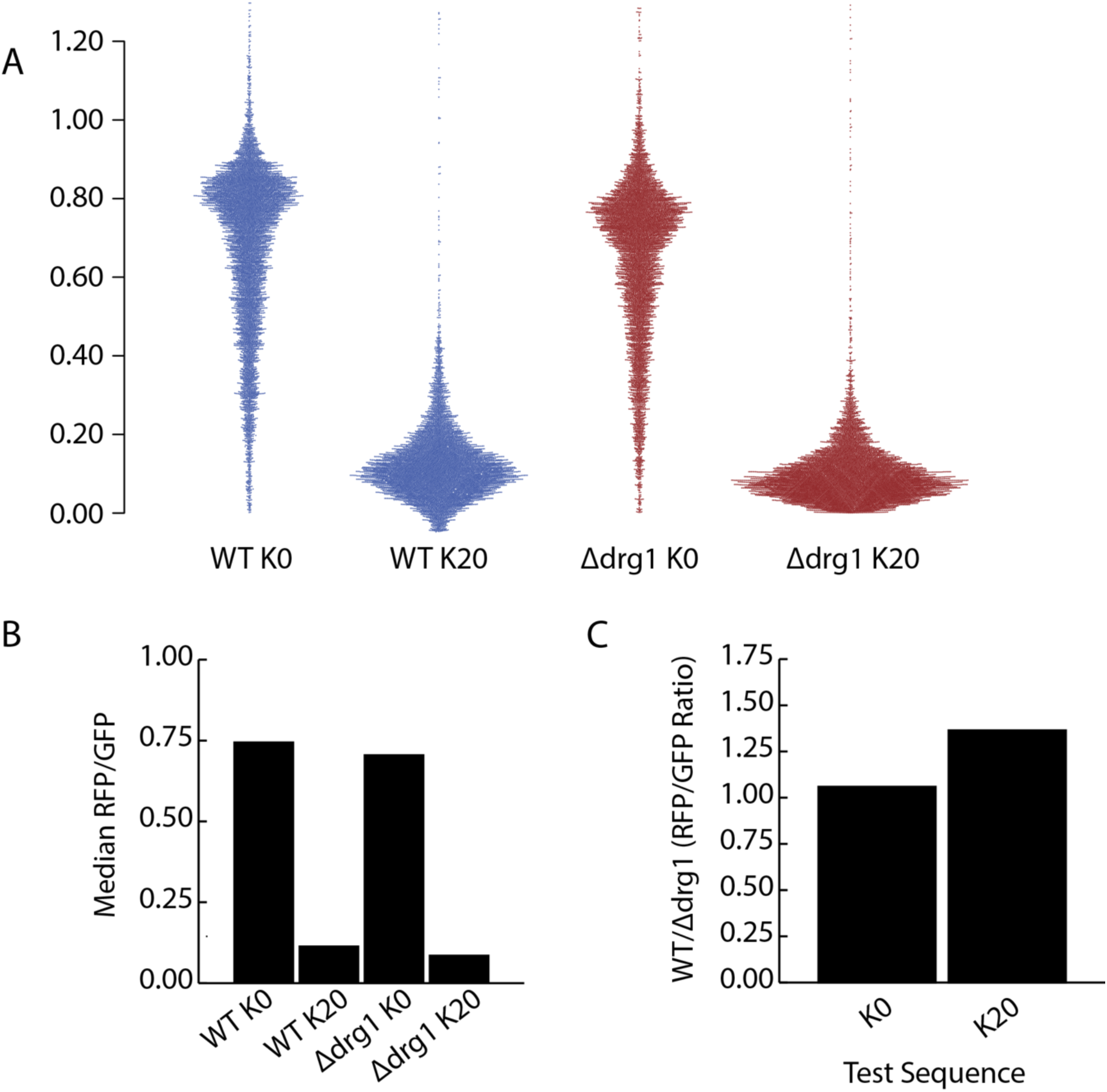
Translation-promoting function of Drg1 is more pronounced in K20 stalling sequences than K0 controls. **A. Beeswarm plot showing the RFP/GFP ratios of approximately 8000 cells per experiment**. WT and Δdrg1 cells were transfected with a dual fluorescent reporter construct encoding a test sequence of either zero lysines (K0) or twenty lysines (K20) and analyzed by flow cytometry. **B. Median RFP/GFP ratios for the indicated cell treatments**. **C. Bar graph showing the ratio of RFP/GFP in WT cells to RFP/GFP in Δdrg1 cells for both K0 and K20 reporter constructs**. The elevated ratio observed for K20 constructs indicates that Drg1 translation-promoting activity is more pronounced at polybasic stalling sequences.

**Supplemental Fig. 6.**
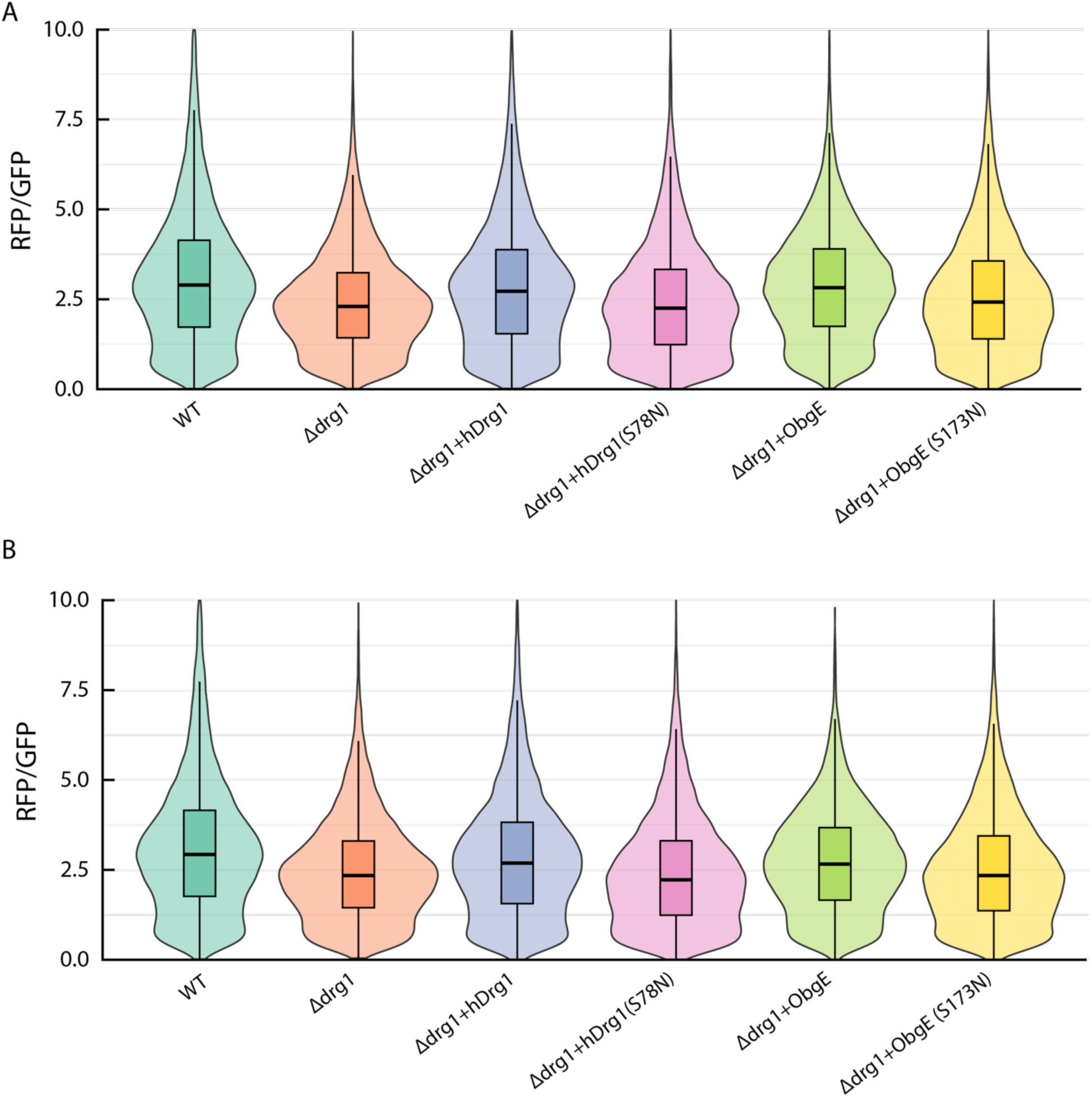
Decreased K20 readthrough in human Δdrg1 cells and its rescue by plasmid-expressed human hDrg1 and bacterial ObgE proteins. Violin plots show RFP/GFP ratios from individual cells co-transfected with the dual-fluorescent reporter and the indicated expression vectors. Medians of the two replicates shown here and the replicate shown in Fig. 2B were used to generate the bar plots and conduct the statistical analyses in Fig. 2C.

**Supplemental Fig. 7.**
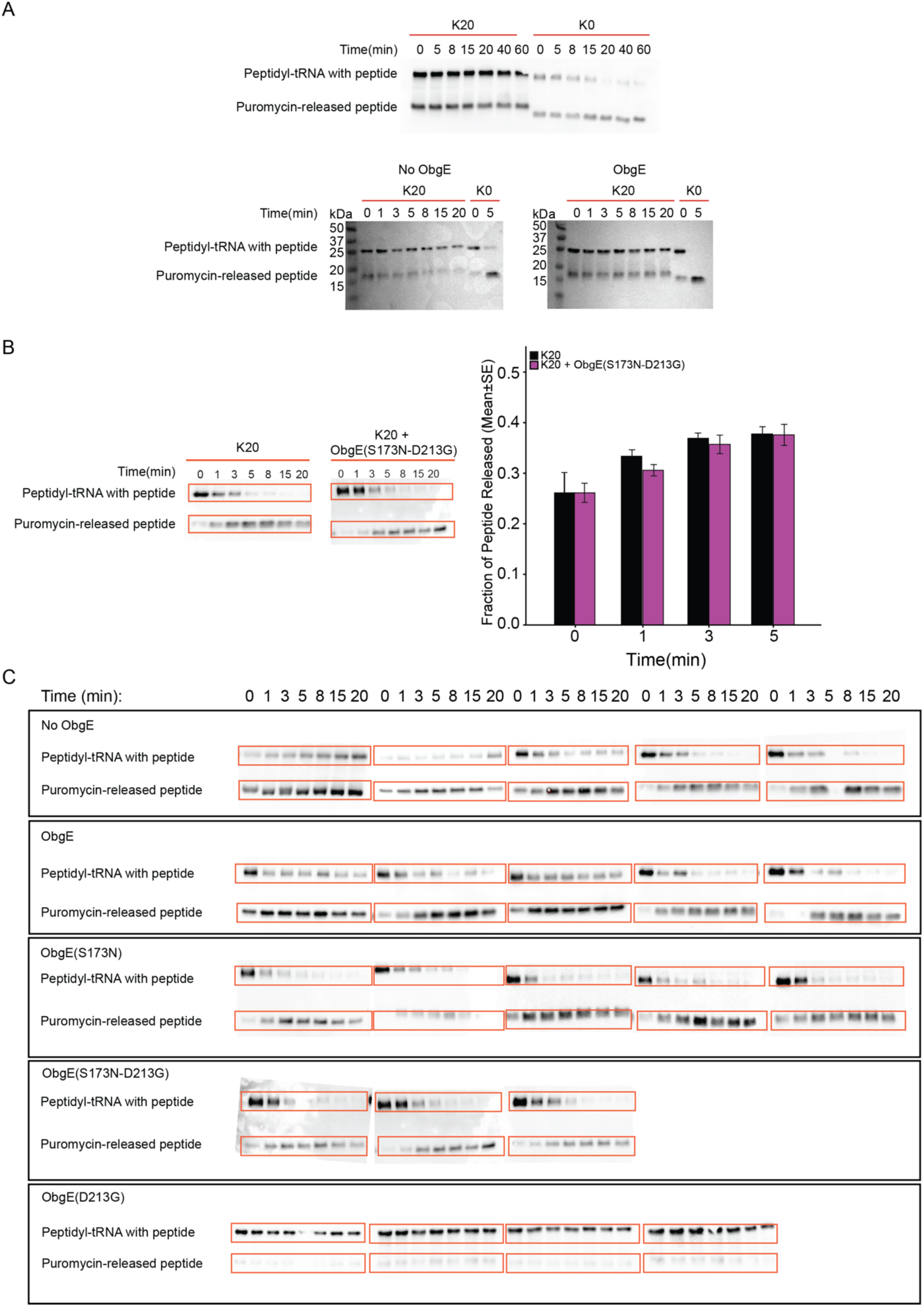
Obg increases puromycin reactivity in the ribosome. **A. ObgE enhances puromycin incorporation in the K20-stalled ribosomes**. Raw *a*FLAG immunoblots were used to show the degree of puromycin incorporation. Representative images are shown for a 0, 5, 8, 15, 20, 40, and 60 min time course used to monitor puromycin reactivity of ribosomes without ObgE that were stalled while translating 20 consecutive AAA codons (K20). Representative images are shown for the same time points of ribosomes translating 3’-truncated mRNA with no consecutive AAA codons (K0). Representative images are shown for a 0, 1, 3, 5, 8, 15, and 20-minute time course used to monitor puromycin reactivity of ribosomes with ObgE that were stalled while translating K20, with the same reaction at K0 (0,5 minutes) as reference. Molecular weight marker is used to indicate the molecular weight differences of the puromycin-released peptide products at K0 and K20 constructs. Unreacted peptidyl tRNA and reacted, puromycylated peptide bands are indicated in the image. **B. ObgE (S173N-D213G) double mutant fails to enhance puromycin incorporation in the stalled ribosome**. Raw *a*FLAG immunoblots were used for puromycin reactivity quantification. Representative images are shown for 0, 1, 3, 5, 8, 15, 20 min time course used to monitor puromycin reactivity of ribosomes that were stalled at K20. Unreacted peptidyl tRNA with nascent peptide attached and reacted puromycin-released peptide bands are indicated in the image. The quantification plot represents the average intensity of the fraction of peptide released, with error bars denoting standard error. See Methods for quantification details. **C. ObgE increases puromycin reactivity in the ribosomes** Raw *a*FLAG immunoblots were used for puromycin reactivity quantification. A minimum of three replicates are shown for 0, 1, 3, 5, 8, 15, 20 min time course used to monitor puromycin reactivity of ribosomes stalled at K20 using ObgE and various ObgE mutants (S173N, D213G, S173N-D213G). Unreacted peptidyl tRNA and reacted puromycylated peptide bands are indicated in red boxes. The figure shows the results from biological replicates. See Methods for quantification details.

**Supplemental Figure 8.**
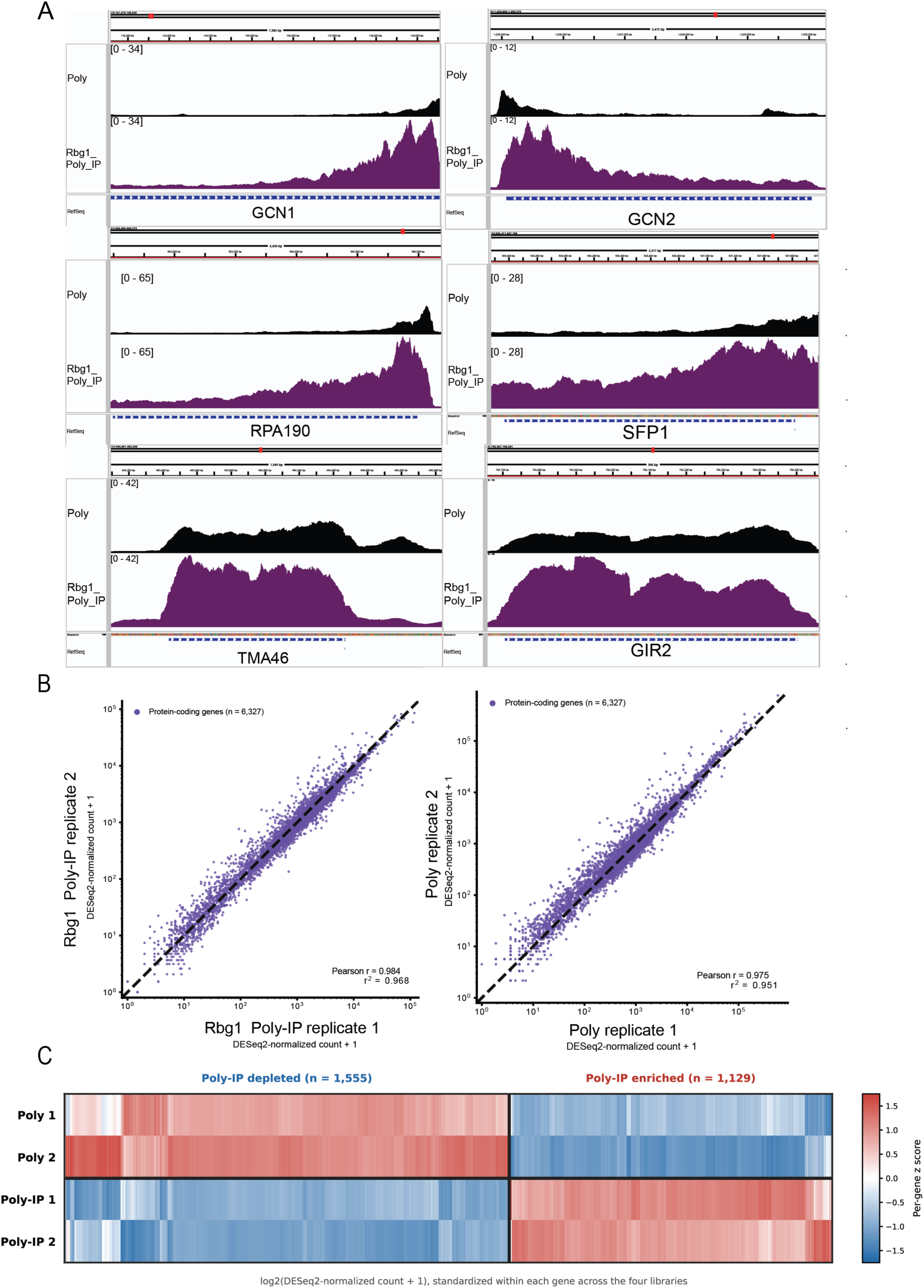
Representative Rbg1 Poly-IP RNA coverage and sequencing-data quality. **A. Representative browser profiles of Rbg1 Poly-IP-enriched transcripts**. Mean RPM-normalized polysome coverage is shown in black and Rbg1 Poly-IP coverage in purple across GCN1, GCN2, RPA190, SFP1, TMA46, and GIR2. GCN1 encodes a ribosome-associated activator of Gcn2, whereas GCN2 encodes the eIF2α kinase that remodels translation during amino-acid limitation and elongation stress. RPA190, the largest RNA polymerase I subunit, supports ribosome biogenesis through 35S pre-rRNA transcription. SFP1 regulates ribosomal-protein and ribosome-biogenesis genes. TMA46 is the established Rbg1 binding and stabilizing partner, while GIR2 is an RWD-domain translation-control factor connecting the Rbg network with Gcn1-dependent regulation. Rbg1 Poly-IP coverage exceeded polysome coverage across all six transcripts. The longer GCN1, GCN2, RPA190, and SFP1 transcripts displayed 3′-end Rbg1 Poly-IP accumulation, whereas TMA46 and GIR2 showed broader coding-region enrichment. Gene models indicate transcript orientation and coding structure. **B. Reproducibility of polysome and Rbg1 Poly-IP libraries**. Scatterplots compare DESeq2-normalized protein-coding counts between biological replicates. Rbg1 Poly-IP replicates showed r = 0.984 and r² = 0.968, whereas polysome replicates showed r = 0.975 and r² = 0.951, across 6,327 genes. Axes show normalized counts plus one on logarithmic scales; dashed lines indicate equal replicate abundance. **C. Expression patterns distinguishing Rbg1 Poly-IP-enriched and Poly-IP-depleted transcripts**. Genes were classified by paired DESeq2 analysis using Benjamini-Hochberg-adjusted p < 0.05 and absolute log_2_ fold change > 1. Heatmap values are log₂(normalized count + 1), standardized within each gene across four libraries. Red indicates above-gene-mean expression, blue indicates below-mean expression, and white indicates values near the mean. Depleted genes showed higher polysome signal, whereas enriched genes showed higher recovery in both Rbg1 Poly-IP replicates.

**Supplemental Fig. 9.**
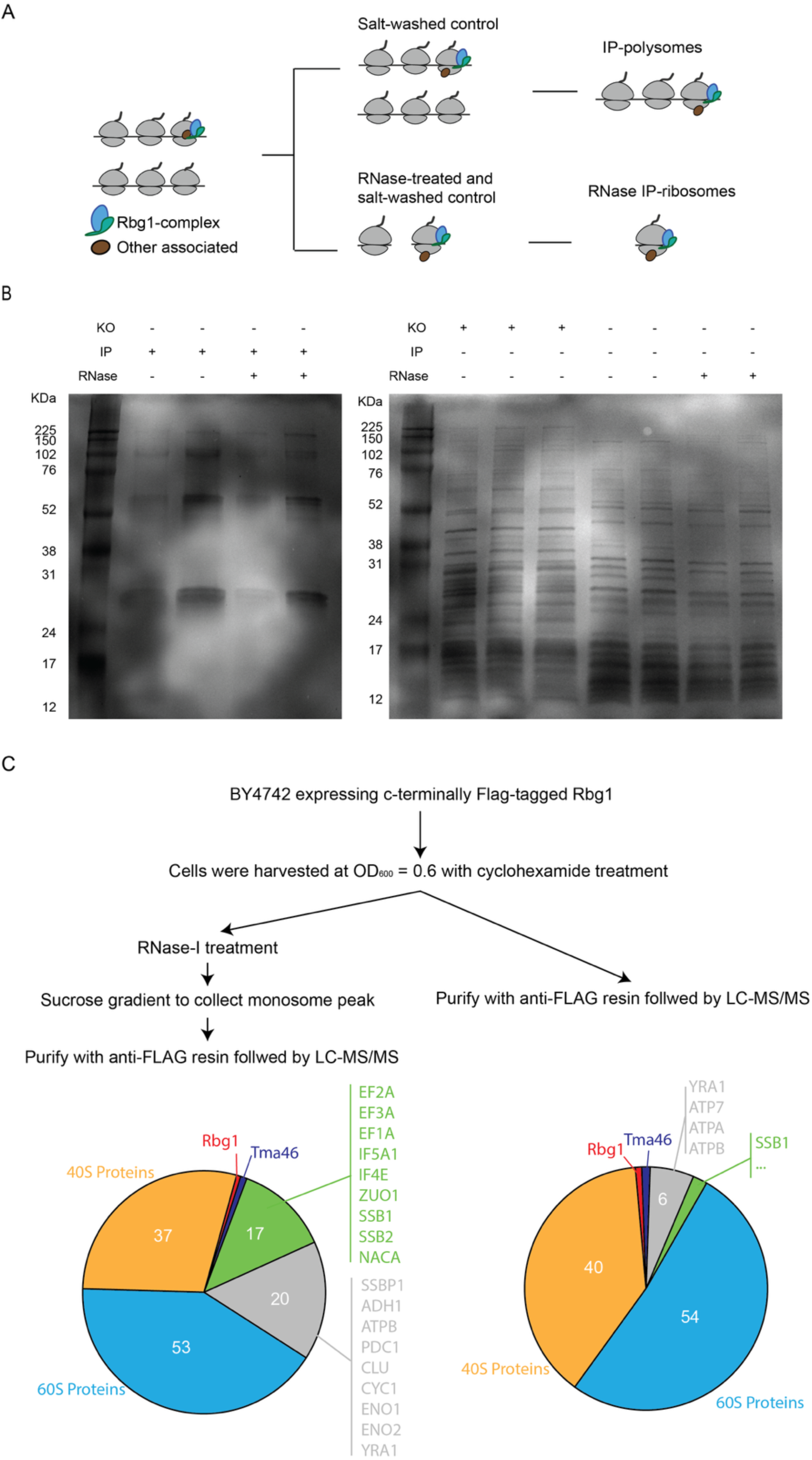
Identification of peptides associated with Rbg1-80S translation complex by IP-LC-MS/MS. **A. Schematic of the IP-LC-MS/MS workflow**. Mid-log phase yeast expressing C-terminally FLAG-tagged Rbg1 (chromosomal) were treated with 100 μg/mL cycloheximide. 80S ribosomes were isolated via 1M sucrose cushion sedimentation, and Rbg1-bound ribosomes were immunoprecipitated using αFLAG affinity gel. Four types of samples were generated: total 80S with additional salt wash of the ribosomal fraction (Salt-washed control) to match the required salt wash for Rbg1-Flag immunoprecipitation (IP), Rbg1-Flag IP of the ribosomal fraction (IP-polysomes), RNase treatment prior to sucrose cushion followed by salt wash (RNase-treated Salt-washed control), and RNase-treated ribosomal fraction followed by Rbg1-Flag IP (RNase_IP). Three biological replicates were used to analyze by LC-MS/MS. **B. SDS–PAGE quality control of ribosome-associated pulldown for mass spectrometry**. Coomassie-stained SDS–PAGE gel showing affinity-purified ribosome-associated complexes used for LC–MS/MS analysis. Samples include pulldown of Rbg1-ribosome complex with or without RNase I treatment. Translation fractions of Δrbg1 cell were also studied to study the cellular response in the absence of Rbg1, and the results are to be reported elsewhere. The bait protein and other co-purifying bands show ribosomal and associated factors. These data provide the input material for mass spectrometry analysis. The gel was stained with Coomassie blue and image using iBright1500 (Thermo). **C. Rbg1 GTPase associates with ribosomes and other translation factors**. Summary of experimental procedure and results on proteins co-immunoprecipitated with Rbg1 upon high-salt wash. Interacting proteins are grouped into the following categories: 60S ribosomal proteins (blue), 40S ribosomal proteins (orange), translation factors and Nascent peptides (green), Tma46 (dark blue), Rbg1 itself (red), and other proteins (grey). Note that Rbg1/Tma46-bound ribosomal complexes can be directly pulled down from the cell and elute was sent to mass spectrometry analysis.

